# The Mycobacterium tuberculosis ESX effector promotes pyroptosis-dependent pathogenicity and dissemination

**DOI:** 10.1101/2025.05.13.653593

**Authors:** Yajie Shen, Yifan He, Anke Chen, Yuanyuan Li, Yuhui Gao, Xuehe Liu, Lu Geng, Menglin Ye, Yuxin Qiu, Lu Zhang, Yicheng Sun, Hua Yang, Jixi Li

## Abstract

Graphic Abstract

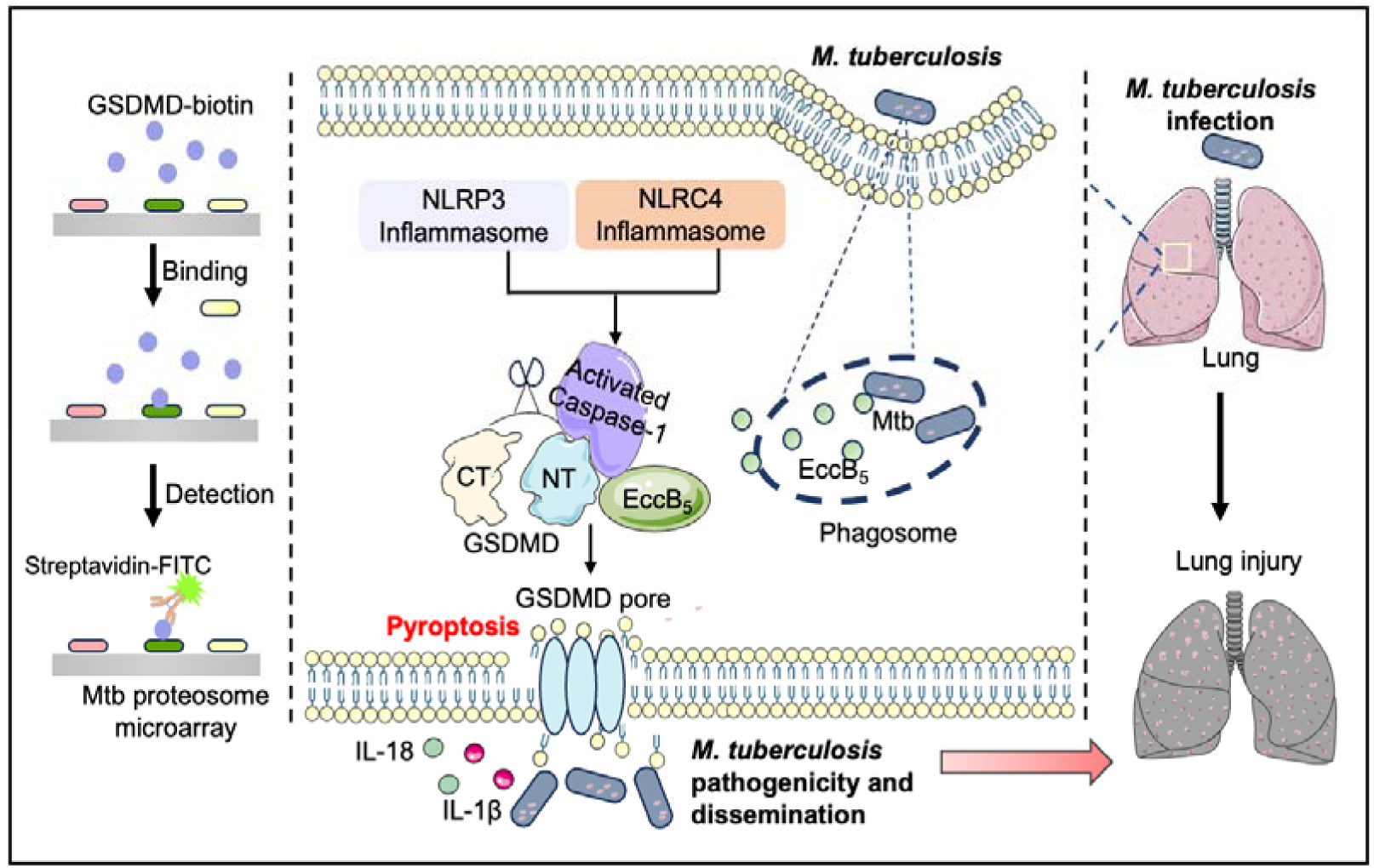

*Mycobacterium tuberculosis* (Mtb) remains a major global health threat, partly due to the extensive cytotoxicity induced during infection. Although GSDMD-mediated excessive pyroptosis promotes pathogen dissemination and tissue damage in tuberculosis, the mechanism remains poorly understood. Here, we identify the effector EccB_5_, a component of the Mtb ESX-5 secretion system, as a key driver of pyroptosis and hyperinflammatory responses. EccB_5_ enhances Mtb virulence by inducing pyroptosis of macrophages, promoting bacterial dissemination and exacerbating lung pathology. Conditional knockdown EccB_5_ increases host cell viability. Mechanistically, EccB_5_ directly interacts with GSDMD, strengthens its association with caspase-1, and facilitates caspase-1-mediated cleavage of GSDMD both *in vitro* and *in vivo*. In summary, our findings uncover a precise mechanism by which Mtb modulates host responses and advances its pathogenicity.

## Introduction

Tuberculosis (TB), caused by *Mycobacterium tuberculosis* (Mtb), remains one of the leading causes of infectious diseases worldwide and continues to pose a significant global public health challenge [1, 2]. Current anti-TB therapies under clinical investigation primarily focus on inhibiting bacterial growth and metabolism. While effective in the short term, such strategies may contribute to the development of drug resistance over prolonged use [3–5]. In contrast, host-directed therapy (HDT) has emerged as a promising alternative that targets key host cellular pathways to enhance immune responses and facilitate pathogen clearance [6–8]. HDT strengthens intrinsic cellular processes to counter infection, emphasizing the need for a deeper understanding of host-Mtb interactions during the immune response [9, 10].

Pyroptosis is a lytic and pro-inflammatory form of programmed cell death characterized by membrane pore formation and release of intracellular inflammatory mediators [11, 12]. This process is primarily driven by the inflammasome-mediated cleavage and activation of gasdermin D (GSDMD), the key executioner of pyroptosis [13–15]. Accumulating evidence highlights the essential role of pyroptosis in controlling Mtb infection, by eliminating intracellular pathogens and activating immune signaling pathways [16, 17]. However, pyroptosis can also promote tissue damage and bacterial dissemination if dysregulated. Pyroptosis can be triggered by phagosome rupture in infected macrophages, allowing bacterial components to access the cytosol and activate inflammasome signaling. Although this response helps contain infection initially, sustained or excessive pyroptosis may compromise host tissue integrity and facilitate Mtb spread [18–20]. Despite growing recognition of the importance of pyroptosis in TB pathogenesis, the mechanisms by which Mtb manipulates host cell pyroptosis remain largely unclear.

In this study, we conducted a comprehensive proteomic screen using an Mtb protein microarray to identify bacterial factors involved in regulating macrophage pyroptosis, and discovered EccB_5_, a protein encoded by the ESX-5 secretion system gene Rv1782c [21], as an inducer of pyroptosis. ESX-5 functions in the induction of interleukin-1β (IL-1β) release during mycobacterial infection [22]. Here, we show that EccB_5_ directly interacts with GSDMD, promoting its cleavage and triggering pyroptotic cell death in macrophages. Conditional knockdown of EeccB_5_ in the virulent H37Rv strain increases host cell survival, whereas overexpression of EccB_5_ in *Mycobacterium smegmatis* activates pyroptosis, reduces bacterial clearance, and exacerbates tissue damage. These results provide novel insights into the regulatory role of Mtb-derived proteins in host cell death and broaden our understanding of the complex interplay between host immunity and Mtb pathogenesis.

## Results

### Mtb EccB_5_ induces macrophage pyroptosis

To identify potential effectors of Mtb involved in the regulation of pyroptosis, we purified recombinant human GSDMD protein and used it to probe an Mtb proteome microarray (Fig. 1A and B). Based on stringent selection criteria (as described in the Experimental Procedures), we identified two GSDMD-interacting proteins: EccB_5_ (encoded by the Rv1782c gene) and Rv2315c (Table S1). To assess their potential functions, we measured the viability of wild type (WT) iBMDM cells and cells overexpressing either EccB_5_ or Rv2315c following stimulation with pyroptosis stimulator LPS/Nigericin or LPS/flagellin. Notably, iBMDM cells overexpressing EccB_5_ exhibited increased sensitivity to LPS/Nigericin-induced pyroptosis compared to Rv2315c-expressing cells (Fig. 1C). Consistently, THP-1 cells stably expressing EccB_5_ were also more susceptible to pyroptotic cell death upon LPS/Nigericin or LPS/flagellin treatment (Fig. 1D). EccB_5_ is a core component of the ESX-5 secretion system [21], which plays a critical role in host-pathogen interactions through its effects on PPE protein secretion, cell wall integrity, and virulence [23, 24]. Due to the essentiality of Rv1782c for Mtb viability, we utilized a CRISPR interference (CRISPRi) strategy [25] to generate an anhydrotetracycline (ATc)-inducible Rv1782c knockdown strain (EccB_5__cKDTet) in the Mtb H37Rv background (Fig. 1E). Upon ATc induction, knockdown of EccB_5_ significantly reduced cell pyroptosis in *Gsdmd*^+/+^ iBMDMs but had no effect in *Gsdmd*^-/-^ iBMDMs (Fig. 1F and G). Also, knockdown of EccB_5_ suppressed the secretion of the key proinflammatory cytokines IL-1β and IL-18, both of which are released upon inflammasome activation in a GSDMD-dependent manner (Fig. 1H and I). Moreover ATc-induced knockdown of EccB_5_ led to a marked reduction in intracellular Mtb survival in *Gsdmd*^+/+^ iBMDMs, whereas no significant change was observed in *Gsdmd*_/_ iBMDMs (Fig. 1J).

**Fig. 1.**
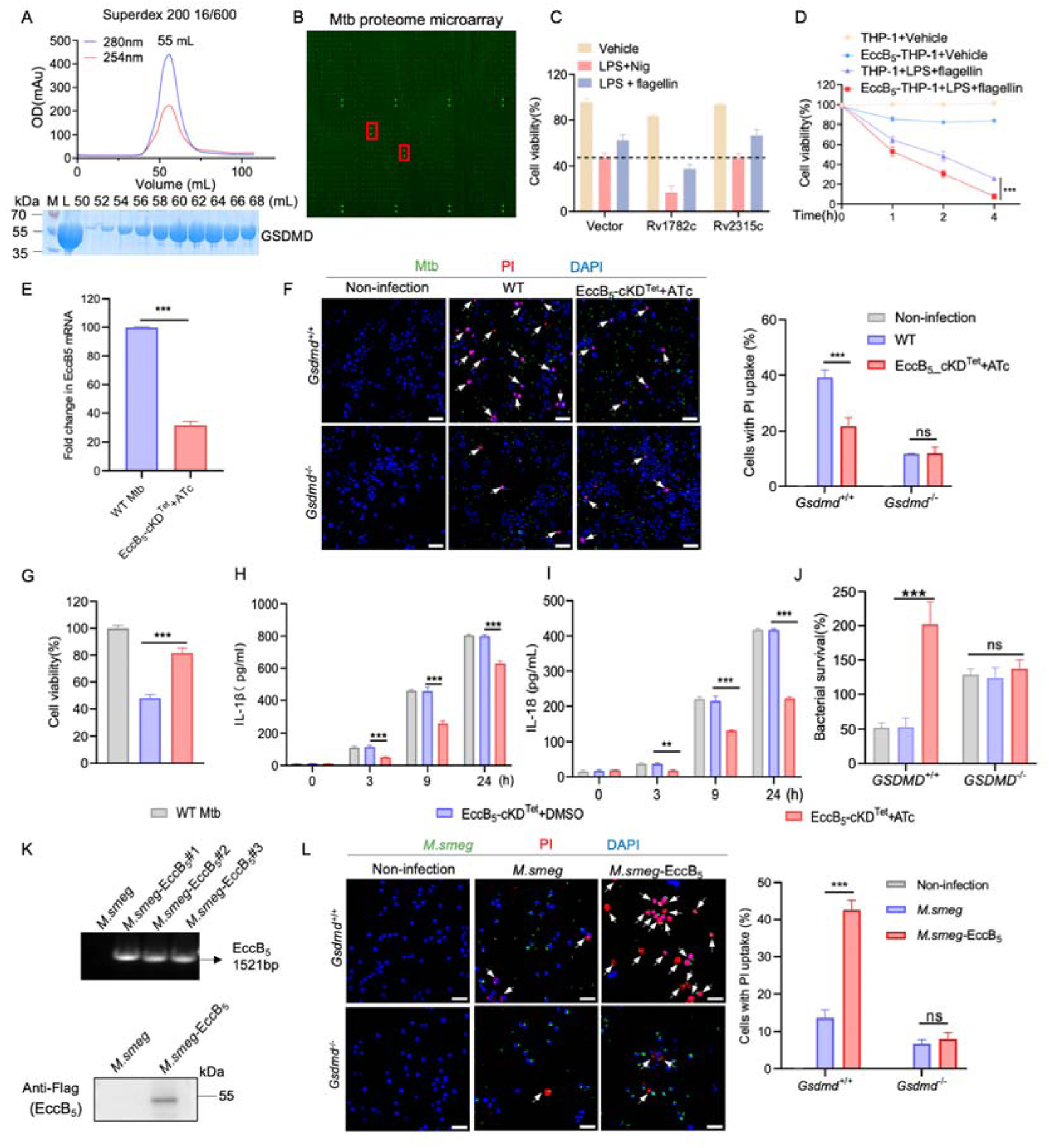
*M. tuberculosis* EccB_5_ induces pyroptosis of macrophages. (**A**) Size-exclusion chromatographic profile and SDS-PAGE result of the GSDMD protein on a Superdex200 16/600 GL column. M: protein marker. (**B**) The full view of a representative tuberculous-associated protein microarray detected with anti-GSDMD signal. Two potential interacting proteins were highlighted with red rectangles. (**C**) Cell viability of iBMDM cells by Cell Counting Kit-8 (CCK-8) assay. WT iBMDM cells and cells stably expressing Rv1782c (EccB_5_-iBMDM) and Rv2315c (Rv2315c-iBMDM) were treated with vehicle, 1 μg/mL LPS, 20 μM nigericin, or 30 μM flagellin for the indicated time periods. **(D)** Cell viability of THP-1 cells by Cell Counting Kit-8 (CCK-8) assay. WT THP-1 cells and THP-1 cells stably expressing EccB_5_ (EccB_5_-THP-1) were treated with vehicle, 1 μg/mL LPS, 20 μM nigericin, or 30 μM flagellin. for the indicated time periods. (**E**) qPCR analysis of Rv1782c mRNA from WT H37Rv and ATc induced Rv1782c knockdown strains (EccB_5__cKDTet). (**F**) Membrane integrity and PI uptake analysis of iBMDMs upon infection with WT H37Rv and ATc induced Rv1782c knockdown strains. The Arrows indicate propidium iodide (PI)-positive cells. Scale bars, 50 μm. (**G**) Analysis for cell cytotoxicity in iBMDMs, in corresponding with (**F**). **(H** and **I)** ELISA of supernatant IL-1β (**F**) and IL-18 (**G**) from iBMDMs. (**J**) Intracellular survival analysis of Mtb in iBMDMs. For (**F**) to (**J**), cells were infected with or without Mtb strains for 24 hours. (**K**) PCR amplification of EccB_5_ encoding sequence from *M. smeg* or *M. smeg*-EccB_5_ strains and western blot analysis of the expression of Flag-tagged EccB_5_ protein in *M. smeg* or *M. smeg*-EccB_5_ strains. (**L**) Membrane integrity and PI uptake analysis of iBMDMs upon infection with *M. smeg* or *M. smeg*-EccB_5_ strains. The Arrows indicate PI-positive cells. Scale bars, 50 μm. Cells were infected with the indicated *M. smeg* strains for 48 hours. Data are presented as mean ± SD of three independent groups. Statistical significance was determined by two-way ANOVA followed by Tukey’s multiple comparisons test. ns, not significant (P > 0.05); P < 0.05; *P < 0.01; **P < 0.001; ***P < 0.0001.

Given that *Mycobacterium smegmatis* (*M. semg*) lacks the Rv1782c gene and is widely used as a nonpathogenic model of Mtb, we generated a recombinant strain expressing H37Rv EccB_5_ (designated *M. smeg*-EccB_5_) (Fig. 1K and S1A). Macrophages were infected with recombinant *M. smegmatis* strains, and EccB_5_ expression was found to enhance bacterial intracellular survival in iBMDM and THP1 cells, respectively (Fig. S1B and C). Immunoblot analysis of subcellular fractions from lysed *M. smeg*-EccB_5_ cells revealed that EccB_5_ was predominantly localized to the cell membrane and cell wall (Fig. S2A). Moreover, treatment of intact *M. smeg*-EccB_5_ cells with proteinase K led to degradation of surface-exposed EccB_5_, further confirming its surface accessibility (Fig. S2B). As conserved homologs of genes neighboring *eccB_5_*in related species are implicated in membrane association or cell wall biosynthesis, EccB_5_ might participate in cell wall-related processes, of which perturbations can significantly alter colony morphology [26]. While colonies of WT *M. smeg* appeared dry, brittle, and wrinkled with irregular margins, *M. smeg*-EccB_5_ formed glossy, smooth, moist, and rounded colonies (Fig. S2C). Notably, expression of EccB_5_ did not affect the overall growth kinetics of *M. smegmatis* (Fig. S2D). However, EccB_5_ overexpression altered colony morphology, delayed biofilm formation, and did not affect bacterial survival under acidic stress conditions (Fig. S2E to H). Functionally, infection of macrophages with *M. smeg*-EccB_5_ resulted in significantly decreased host cell viability, accompanied by increased lactate dehydrogenase (LDH) release, propidium iodide (PI) uptake, and interleukin-1β (IL-1β) secretion, relative to infection with WT *M. smeg* (Fig. 1K and S3A-C), indicating enhanced induction of pyroptotic cell death.

Next, the recombinant EccB_5_ protein was introduced into RAW264.7 cells via electroporation, which induced lytic cell death characterized by ballooning morphology (Fig. S4A and B), a hallmark of pyroptosis [27]. In contrast, direct addition of EccB_5_ to the culture medium had no effect on cell viability or morphology (Fig. S4B and C), suggesting that EccB_5_ requires cytosolic localization and does not act via a cell surface receptor. Furthermore, EccB_5_ induced macrophage death in a time-dependent manner (Fig. S4D and E). The presence of polymyxin B, an LPS neutralizer [28], did not affect EccB_5_-induced cell death across a range of concentrations (Fig. S4F and G), indicating that the cytotoxic effects of EccB_5_ are independent of LPS stimulation. Taken together, these findings identify EccB_5_ as a cytosolic trigger of GSDMD-dependent pyroptosis, promoting macrophage death and proinflammatory cytokine release.

### Mtb EccB_5_ directly interacts with GSDMD

To validate the interaction between GSDMD and EccB_5_, GFP tagged EccB_5_ was transfected into HEK293T cells, and immunoprecipitation with anti-GFP antibody identified GSDMD as one of the top hints by the LC-MS/MS method (Fig. S5A and table S2). Next, we performed co-immunoprecipitation (co-IP) using HA-tagged human GSDMD and Flag-tagged EccB_5_. As expected, Flag-EccB_5_ successfully co-immunoprecipitated with HA-GSDMD (Fig. 2A). A reciprocal co-IP experiment further supported this binding capability (Fig. 2B). Moreover, exogenous co-IP assays revealed that EccB_5_ could associate with both human and murine GSDMD (Fig. 2C and D), and endogenous co-IP in THP-1 cells and iBMDMs confirmed this interaction under physiological conditions, which was further enhanced upon inflammasome activation (Fig. 2E and F). Interestingly, EccB_5_ interacted not only with full-length GSDMD but also with its individual domains—the N-terminal fragment (residues 1–144 or 1–176) and the C-terminal region—as demonstrated by co-IP assays (Fig. 2G and S5B), indicating that EccB_5_ can bind both GSDMD-N and GSDMD-C.

**Fig. 2.**
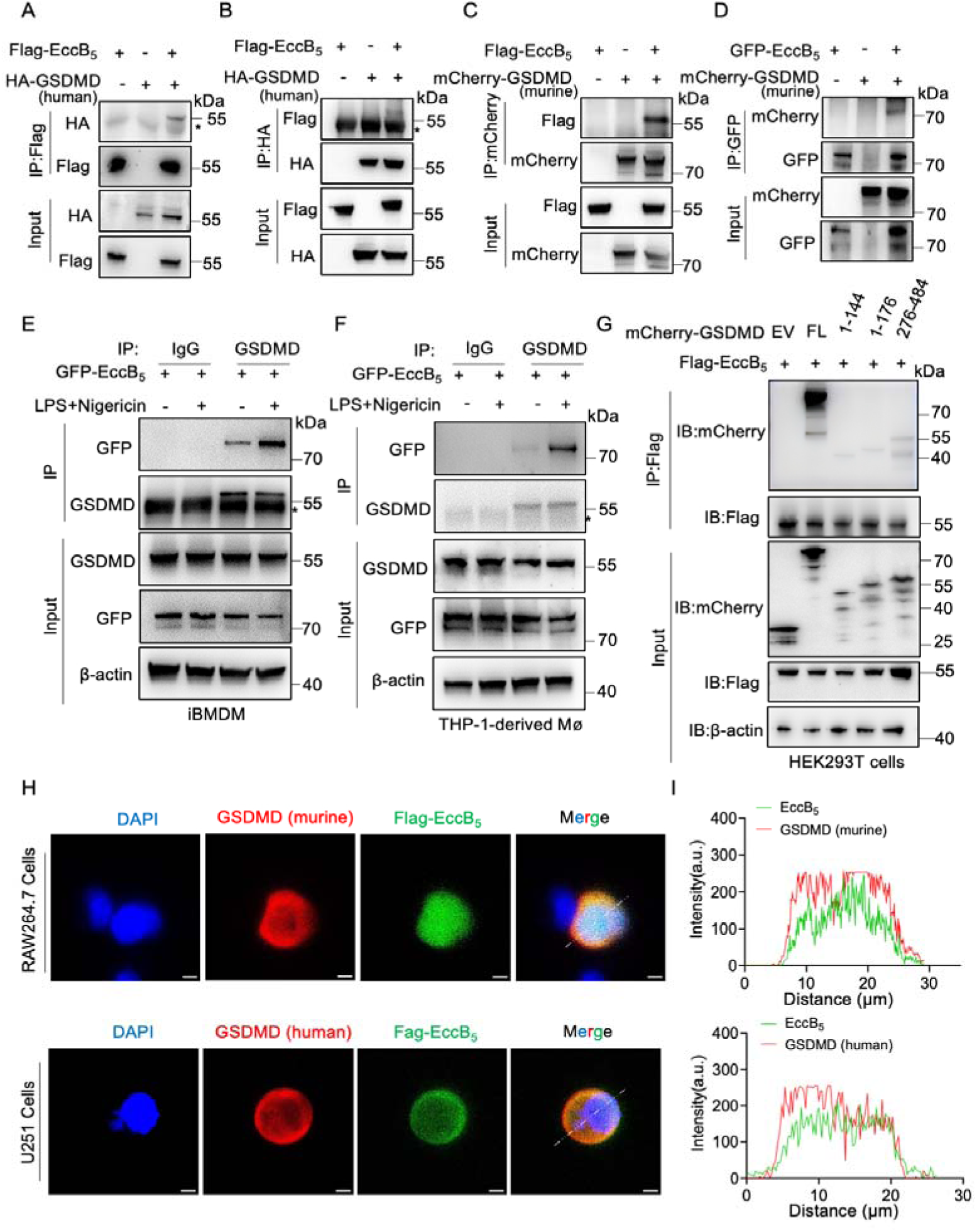
*M. tuberculosis* EccB_5_ directly interacts with GSDMD. **(A to D)** Coimmunoprecipitation (Co-IP) analysis of Flag-EccB_5_ or GFP-EccB_5_ with human HA-GSDMD or murine mCherry-GSDMD in HEK293T cells. Asterisk (*) indicates IgG binding. (**E** and **F**) iBMDMs (**E**) or THP-1 cells (**F**) stably expressing GFP-tagged EccB_5_ were pretreated with 1 μg/mL LPS for 2 h, followed by 20 μM nigericin for an additional 2 h. Cell lysates were immunoprecipitated with anti-Flag or control IgG, and both inputs and precipitates were analyzed by immunoblotting. (**G**) HEK293T cells were transfected with GSDMD-mCherry or its mutants and Flag-EccB_5_. Cell lysates were immunoprecipitated with anti-Flag antibody and then immunoblotted with indicated antibodies antibody. EV: empty vector. (**H**) RAW264.7 cells and U251 cells were transfected with Flag-EccB_5_ plasmid. The cells were processed for confocal microscopy with anti-Flag antibody (EccB_5_) and anti-GSDMD antibody. Nuclei were stained with DAPI. Scale bars, 5 μm. (**I**) Normalized intensity profiles are drawn from the white lines in (**H**) and show the relative pixel intensity along the line with regards to the distance and fluorescence wavelength (red, Cy3; green, Alexa 488).

To assess subcellular localization, confocal microscopy of HEK293T cells co-expressing Flag-EccB_5_ and HA-GSDMD showed cytoplasmic co-distribution of the two proteins, consistent across both human and murine GSDMD in U251 and RAW264.7 cells, respectively (Fig. 2H and I). Additionally, ectopic expression of EccB_5_ in HEK293T cells also demonstrated co-localization with endogenous GSDMD (Fig. S5C and D), further validating the physiological relevance of this interaction.

EccB_5_ is an essential structural component of the ESX-5 type VII secretion system, which is typically surface-exposed on the bacterial cell wall [29]. The cell fractionation analysis of *M. smeg*-EccB_5_ showed that EccB_5_ was predominantly localized in the cell wall fractions (Fig. S2A), similar to the known surface protein Ag85B, which served as a control [30]. Next, we examined whether EccB_5_ could mediate direct binding of GSDMD to the mycobacterial surface. After incubating recombinant GSDMD with WT *M. smegmatis* or *M. smeg*-EccB_5_, we observed enhanced binding of GSDMD to the EccB_5_-expressing strain (Fig. S5E). Furthermore, treating *M. smeg*-EccB_5_ cells with an exposure to proteinase K before GSDMD incubation resulted in degradation of surface-exposed EccB_5_ and Ag85B, and this degradation significantly reduced GSDMD binding (Fig. S5E). Together, these findings confirm that *M. tuberculosis* EccB_5_ interacts directly with GSDMD, both in the host cytoplasm and on the bacterial surface.

### The EccB_5_ C terminus and R406 are the critical sites for GSDMD binding

To further elucidate the binding interface between EccB_5_ and GSDMD, we employed AlphaFold3-based structural modeling to predict the complex structure of the two proteins (Fig. 3A). The result showed that the C-terminus of EccB_5_ bound with GSDMD, and three interactive pairs of residues were identified in the interface (Arg406-Glu410, Glu434-Gly429, and Trp437-Leu427). Next, a series of GFP-tagged EccB_5_ truncation mutants were designed to identify the specific domain responsible for GSDMD binding. Co-immunoprecipitation assays revealed that the EccB_5_ ^Δ400–506^ mutant, which lacks the C-terminal region (residues 400–506), lost the ability to bind GSDMD (Fig. 3B), indicating that this C-terminal segment is essential for the interaction.

**Fig. 3:**
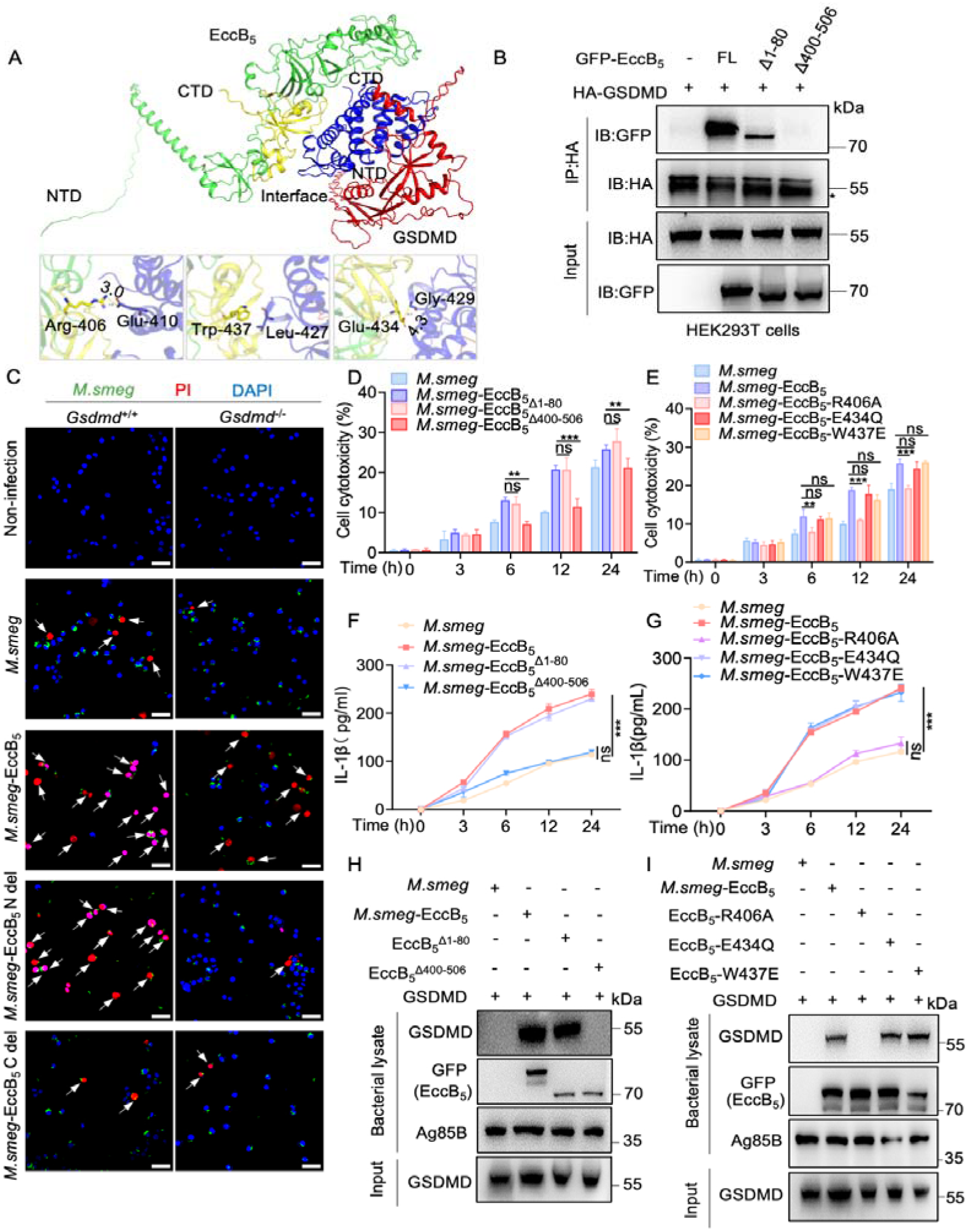
EccB_5_ triggers pyroptosis in a GSDMD-dependent manner. (**A**) The predicted structure of the EccB_5_-GSDMD complex generated using AlphaFold3. Three pairs of interactive residues between the complex are shown in stick models with Pymol software. (**B)** Co-immunoprecipitation and western blotting analysis showing the binding of GFP-EccB_5_ and its variants with HA-GSDMD in HEK293T cells. (**C**) Membrane integrity analysis of iBMDMs upon infection with *M. smeg* or *M. smeg*-EccB_5_ and its mutant strains. The Arrows indicate PI-positive cells. Scale bars, 50 μm. Cells were infected with the indicated *M. smeg* strains for 48 hours. (**D** to **G**) iBMDMs infected with *M. smeg* and *M. smeg*-EccB_5_/EccB ^Δ1-80^/EccB ^Δ400-506^ (**D**) or *M. smeg*-EccB_5_ / EccB_5_-R406A/ EccB_5_-E434Q/ EccB_5_-W437Q (**E**) were evaluated by LDH and ELISA assay. (**H** and **I**) Immunoblot analysis of whole bacterial lysates from the indicated *M. smegmatis* strains and their variants after incubation with GSDMD at 4 °C for 4 hours. Data are presented as mean ± SD of three independent groups. Statistical significance was determined by two-way ANOVA followed by Tukey’s multiple comparisons test. ns, not significant (P > 0.05); P < 0.05; *P < 0.01; **P < 0.001; ***P < 0.0001.

Next, we infected macrophages with WT *M. smegmatis* and various *M. smeg*-EccB_5_ mutants, including EccB_5_ ^Δ400–506^ and a point mutant EccB_5_-R406A. Compared to the WT *M. smeg*-EccB_5_ strain, both mutants showed significantly reduced LDH release, higher cell viability, and decreased PI uptake—hallmarks of reduced pyroptosis (Fig. 3C to E). Consistently, the secretion of IL-1β and IL-18 by macrophages infected with *M. smeg*-EccB_5_ ^Δ400–506^ or *M. smeg*-EccB_5_-R406A was also significantly reduced in a time-dependent manner (Fig. 3F and G), whereas mutants EccB_5_-E434Q and EccB_5_-W437E did not show any difference with *M. smeg*-EccB_5_, further supporting a loss of pro-pyroptotic activity.

To test the binding capability of these mutants, we incubated recombinant GSDMD protein with WT *M. smegmatis* or the EccB_5_ mutant strains. Notably, GSDMD binding to both *M. smeg*-EccB_5_ ^Δ400–506^ and *M. smeg*-EccB_5_-R406A was markedly diminished (Fig. 3H and I), confirming that the C-terminal domain of EccB_5_—particularly the R406 residue—is critical for interaction with GSDMD. Together, these results identify R406 within the C-terminal region of EccB_5_ as a key functional residue required for GSDMD binding and subsequent induction of pyroptosis.

### EccB_5_ triggers macrophage pyroptosis in a GSDMD-dependent manner

To determine whether EccB_5_-induced pyroptosis in macrophages is dependent on GSDMD, we generated *Gsdmd*-knockout iBMDM cells using CRISPR-Cas9 gene editing and exogenously expressed Mtb EccB_5_ in both *Gsdmd*^+/+^ and *Gsdmd*^-/-^ iBMDMs to assess its pro-pyroptotic effect (Fig. 4A) [31]. While EccB_5_ electroporation triggered robust lytic cell death in wild-type cells, GSDMD-deficient macrophages were highly resistant to EccB_5_-induced pyroptosis (Fig. 4B and C), demonstrating a clear dependency on GSDMD for EccB_5_-mediated cytotoxicity.

**Fig. 4.**
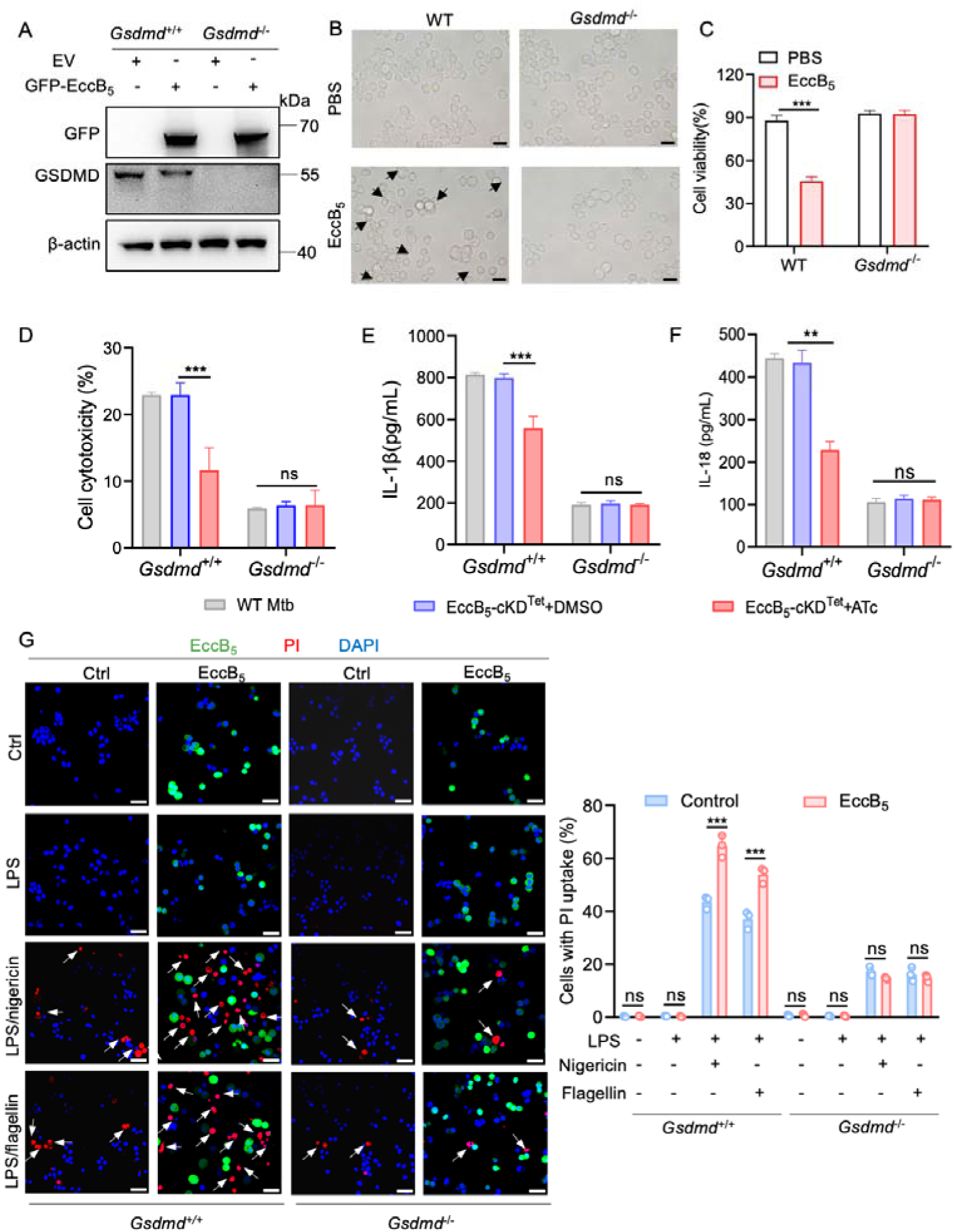
EccB_5_ triggers pyroptosis in a GSDMD-dependent manner. (**A**) Immunoblot analysis of GFP (EccB_5_) and GSDMD in *Gsdmd*^+/+^ and *Gsdmd*^-/-^ iBMDMs stably expressing EccB_5_ or a control vector. (**B** and **C**) WT and *Gsdmd*^-/-^ iBMDMs were transfected with recombinant EccB_5_ or luntreated (PBS) via electroporation for 1 hour. Cell morphology was observed by microscopy (**B**), and cell viability was assessed by CellTiter-Glo luminescent assay (**C**). Scale bar, 20 μm. (**D**) Assessment of cell cytotoxicity in *Gsdmd*^+/+^ and *Gsdmd*^-/-^ iBMDMs infected with Mtb strains. **(E** and **F**) ELISA of supernatant IL-1β (**E**) and IL-18 (**F**) from *Gsdmd*^+/+^ and *Gsdmd*^-/-^ iBMDMs infected with Mtb strains. For **D-F**, cells were infected with or without Mtb H37Rv or ATc induced Rv1782c knockdown strains for 24 hours. (**G**) Membrane integrity analysis in WT and *Gsdmd*^-/-^ iBMDMs stably expressing Mtb EccB_5_, followed by stimulation with LPS/Nigericin or LPS/flagellin. Left, representative fluorescence images showing propidium iodide (PI)-positive cells (arrows). Scale bars, 40 μm. Right, quantification of PI uptake; approximately 100 cells were analyzed. Data are presented as mean ± SD of three independent groups. Statistical significance was determined by two-way ANOVA followed by Tukey’s multiple comparisons test. ns, not significant (P > 0.05); P < 0.05; *P < 0.01; **P < 0.001; ***P < 0.0001.

We next assessed the impact of EccB_5_ knockdown during Mtb infection. Infection with the EccB_5__cKDTet strain led to reduced cell death in WT iBMDMs, whereas no difference in viability was observed between EccB_5__cKDTet and WT H37Rv infections in *Gsdmd*^-/-^ iBMDMs (Fig. 4D). Similarly, IL-1β secretion was unaffected by EccB_5_ knockdown in *Gsdmd*^-/-^ macrophages (Fig. 4E and F). These results suggest that EccB_5_’s effects on pyroptosis and cytokine release are entirely dependent on GSDMD. Consistent findings were observed when infection of *Gsdmd*^-/-^ iBMDMs with either *M. smeg*-EccB_5_ or WT *M. smegmatis* showed no difference in cytotoxicity or inflammatory responses (Fig. S6A and B), further supporting the role of GSDMD in mediating EccB_5_’s effects.

Next, to evaluate GSDMD dependency under inflammasome-activating conditions, we generated WT and *Gsdmd*^-/-^ iBMDMs stably expressing Mtb EccB_5_, followed by stimulation with LPS/Nigericin or LPS/flagellin. In this context, we observed no significant differences in LDH release, cell viability, or IL-1β secretion between WT and *Gsdmd*^-/-^ cells (Fig. 4G, and fig. S6C to F), indicating that GSDMD is essential for EccB_5_-induced pyroptosis, particularly under inflammasome-activating conditions. These data support the notion that EccB_5_ induces GSDMD-dependent pyroptosis to facilitate Mtb intracellular survival.

### EccB_5_ promotes GSDMD cleavage

To further assess the functional consequences of EccB_5_ on GSDMD activation, we compared caspase-1 activation, GSDMD cleavage, and IL-1β secretion in iBMDMs infected with either WT H37Rv or EccB_5__cKDTet knockdown Mtb. Infection with EccB_5__cKDTet resulted in reduced GSDMD-N fragment (35 kDa) formation and significantly decreased IL-1β release compared to WT H37Rv (Fig. 5A). In contrast, compared to WT *M. smegmatis*, infection with *M. smeg*-EccB_5_ led to markedly increased cleavage of full-length GSDMD into its active GSDMD-N and C-terminal (25_kDa) fragments, accompanied by elevated IL-1β secretion in both iBMDMs (Fig. 5B) and human THP-1-derived macrophages (Fig. 5C). Notably, no cleavage of caspase-4/5/11 or caspase-3/GSDME was observed in these experiments (Fig. S6A to D), further confirming that EccB_5_-mediated pyroptosis occurs independently of noncanonical pyroptotic or apoptotic pathway. Moreover, upon LPS/Nigericin stimulation, iBMDMs and THP-1 macrophages stably expressing Mtb EccB_5_ exhibited significantly higher levels of cleaved caspase-1 (p20) and GSDMD-N fragments (Fig. 5D and E), accompanied by elevated IL-1β secretion (Fig. 4H and fig. S6E and F).

**Fig. 5.**
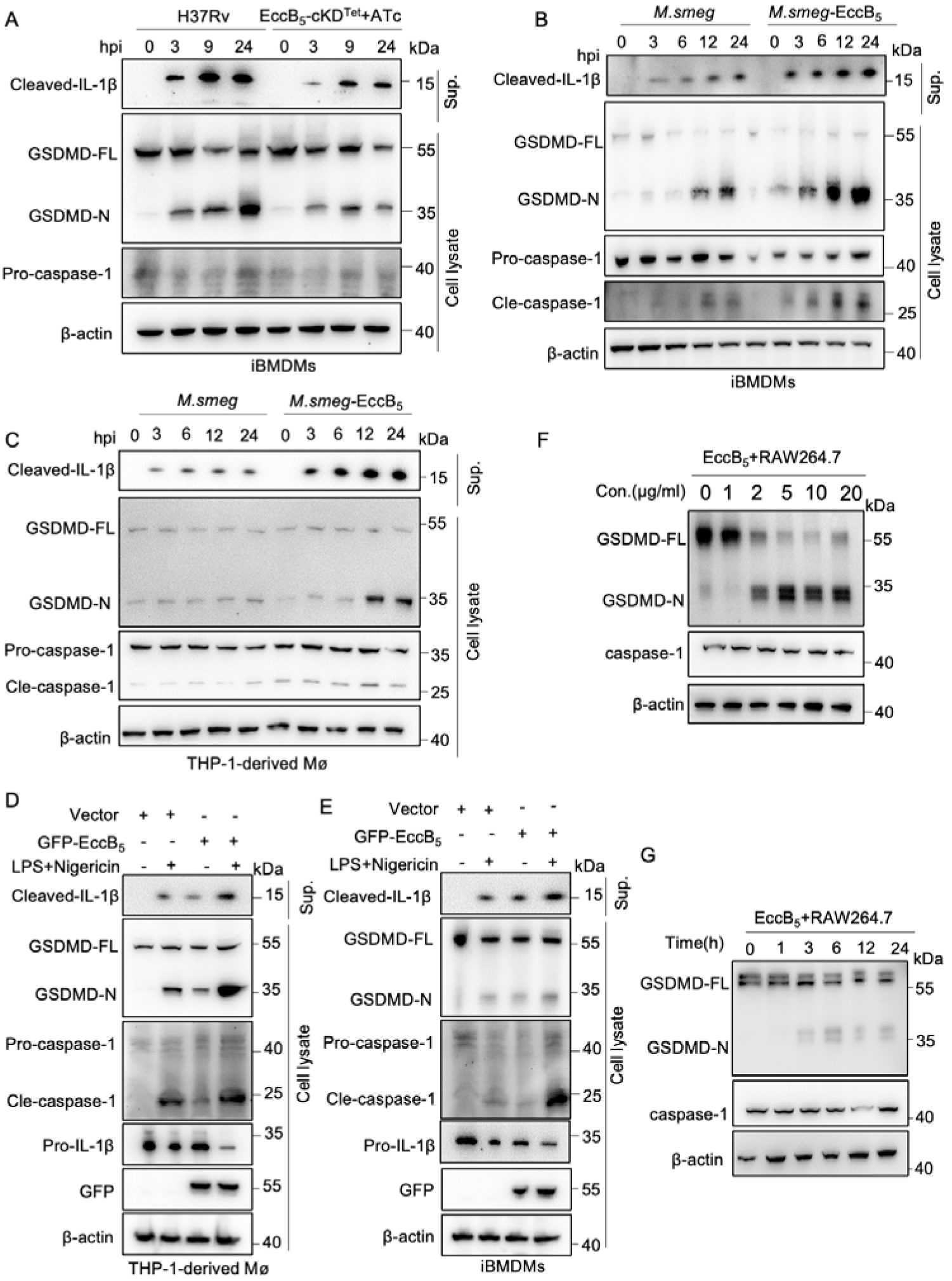
EccB_5_ promotes GSDMD cleavage. (**A** and **B**) iBMDMs were infected with H37Rv, EccB_5__cKDTet, *M. smegmatis*, or *M. smegmatis*–EccB_5_, respectively. Supernatants and cell pellets were collected and analyzed by immunoblotting (WB) with antibodies against the indicated proteins to assess caspase-1, GSDMD cleavage, and IL-1β maturation. (**C**) THP-1 cells infected with *M. smegmatis* or *M. smegmatis*–EccB_5_. Supernatants and cell pellets were collected and immunoblotted with antibodies against the indicated proteins. (**D** and **E**) THP-1 cells or iBMDMs stably expressing EccB_5_ or a control vector were pretreated with 1 μg/mL LPS for 1 h, followed by stimulation with 20 μM nigericin for 2 h. Supernatants and cell pellets were collected and analyzed by immunoblotting with antibodies against the indicated proteins. (**F** and **G**) RAW264.7 cells were electroporated with recombinant EccB_5_ protein at various concentrations (**F**) or electroporated with 2 μg/mL EccB_5_ for different durations (**G**) as indicated. Cell pellets were collected and immunoblotted with antibodies against the indicated proteins.

To investigate whether Mtb EccB_5_ influences GSDMD processing, we electroporated the purified EccB_5_ protein into RAW264.7 macrophages for varying time periods and at different concentrations. Immunoblot analysis revealed that full-length GSDMD was efficiently cleaved into its N-terminal and C-terminal fragments following EccB_5_ treatment (Fig. 5F and G), indicating activation of GSDMD-dependent pyroptosis. In contrast, cleavage of caspase-3 or GSDME was not detected when RAW264.7 cells or THP-1-derived macrophages were infected with the *M. smeg*-EccB_5_ strain (Fig. S7A and B), suggesting that EccB_5_-induced cell death is not mediated through the caspase-3–GSDME axis in mouse macrophages. Moreover, no cleavage differences of caspase-4/5/11 were detected when RAW264.7 cells or THP-1-derived macrophages were infected with the *M. smeg*-EccB_5_ strain (Fig. S7C and D). Also, cleavage of caspase-3 or GSDME was not detected when RAW264.7 cells were electroporated with recombinant EccB5 protein at various concentrations and different durations (Fig. S7E and F). Collectively, these results demonstrate that EccB_5_ enhances GSDMD cleavage and pyroptosis through caspase-1 activation in both murine and human macrophages, underscoring its role as a potent regulator of inflammasome-mediated cell death.

### Interaction of GSDMD with caspase-1 is promoted by EccB_5_

Given the interaction between EccB_5_ and GSDMD, we next sought to determine whether EccB_5_ also modulates components of the inflammasome complex. Co-immunoprecipitation experiments in HEK293T cells revealed that EccB_5_ and caspase-1 could associate with each other in the presence of GSDMD (Fig. 6A and B), raising the possibility that EccB_5_, GSDMD, and caspase-1 form a ternary complex. Notably, EccB_5_ enhanced the interaction between caspase-1 and GSDMD in a dose-dependent manner, as shown by co-IP assays in both HEK293T and RAW264.7 cells (Fig. 6C and D). Thus, while EccB_5_ facilitates the assembly or stabilization of a GSDMD–caspase-1 complex, potentially promoting efficient GSDMD cleavage and pyroptosis.

**Fig. 6.**
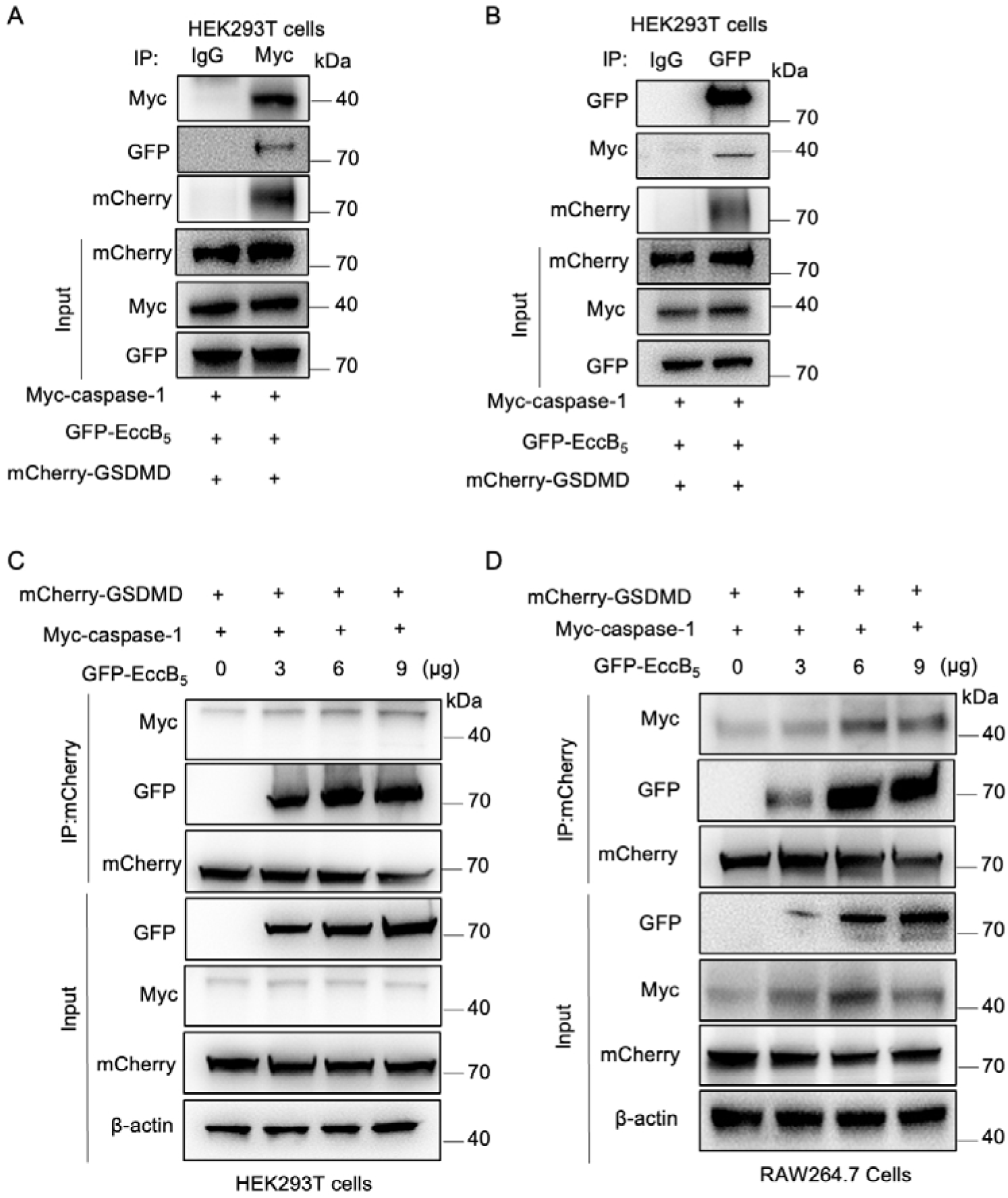
Interaction of GSDMD with caspase-1 is promoted by EccB_5_. (**A** and **B**) HEK293T cells were co-transfected with GFP-EccB_5_, Myc-caspase-1, and mCherry-GSDMD plasmids for 24 h. Cell lysates were subjected to immunoprecipitation (IP) using anti-Myc (**A**) or anti-GFP (**B**) antibodies, followed by immunoblotting with anti-Myc, anti-GFP, and anti-mCherry antibodies. (**C** and **D**) HEK293T (**C**) or RAW264.7 (**D**) cells were co-transfected with varying concentrations of GFP-EccB_5_ together with Myc-caspase-1 and mCherry-GSDMD for 24 h. Cell lysates were immunoprecipitated using an anti-mCherry antibody and analyzed by immunoblotting with anti-Myc, anti-GFP, and anti-mCherry antibodies. 40 μg of total cell lysate were used as input controls.

### EccB_5_ activates host pyroptosis to decrease mycobacterial clearance

To evaluate the role of Mtb EccB_5_ in GSDMD-dependent host immunity, we employed an acute intravenous infection model using *M. smegmatis* and recombinant *M. smeg*-EccB_5_ strains in C57BL/6 mice. Mice were injected with bacteria at 1 × 10_ CFUs per mouse, and lung tissues were assessed at various time points post-infection. At 2 and 6 days post-infection, mice infected with *M. smeg*-EccB_5_ displayed marked histopathological changes in the lungs compared to those infected with wild-type *M. smeg*matis (Fig. 7A). In *Gsdmd*^+/+^ mice, *M. smeg*-EccB_5_ triggered extensive inflammatory cell infiltration, loss of alveolar structure, and progressive tissue destruction. By day 6, histological analysis of *Gsdmd*^+/+^ lungs revealed large areas of necrosis, collapsed alveolar spaces, and extensive tissue disruption in the *M. smeg*-EccB_5_ group, whereas *Gsdmd*^-/-^ mice retained relatively intact lung structure with minimal damage (Fig. 7A and B). In contrast, *Gsdmd*^-/-^ mice showed significantly attenuated pulmonary inflammation and preserved alveolar architecture under the same infection conditions, indicating a critical role for GSDMD in mediating tissue pathology. In parallel, bacterial burden assays demonstrated that infection with *M. smeg*-EccB_5_ resulted in significantly elevated colony-forming units (CFUs) in both the lungs and spleens of *Gsdmd*^+/+^ mice at 2 and 6 days post-infection, as compared to infection with wild-type *M. smegmatis* (Fig. 7C and D). This increase in bacterial burden was associated with excessive pyroptosis triggered by GSDMD activation. This increase in bacterial burden was not observed in *Gsdmd*^-/-^ mice. These results demonstrate that EccB_5_-mediated activation of GSDMD-dependent pyroptosis promotes pulmonary tissue injury and facilitates bacterial expansion and systemic spread. Furthermore, IL-1β and IL-18 levels were significantly elevated in both the serum and lung tissue of *Gsdmd*^+/+^mice infected with *M. smeg*-EccB_5_ at 2 and 6 days post-infection, compared to WT *M. smegmatis* (Fig. 7E to H). These cytokine increases were abolished in *Gsdmd*^-/-^ mice, demonstrating a strict dependency on GSDMD for the proinflammatory effects of EccB_5_. Together, EccB_5_ promotes pyroptosis by targeting GSDMD, leading to lung pathology, elevated inflammatory cytokines, increased bacterial burden, and contributing to Mtb pathogenesis and dissemination *in vivo*.

**Fig. 7.**
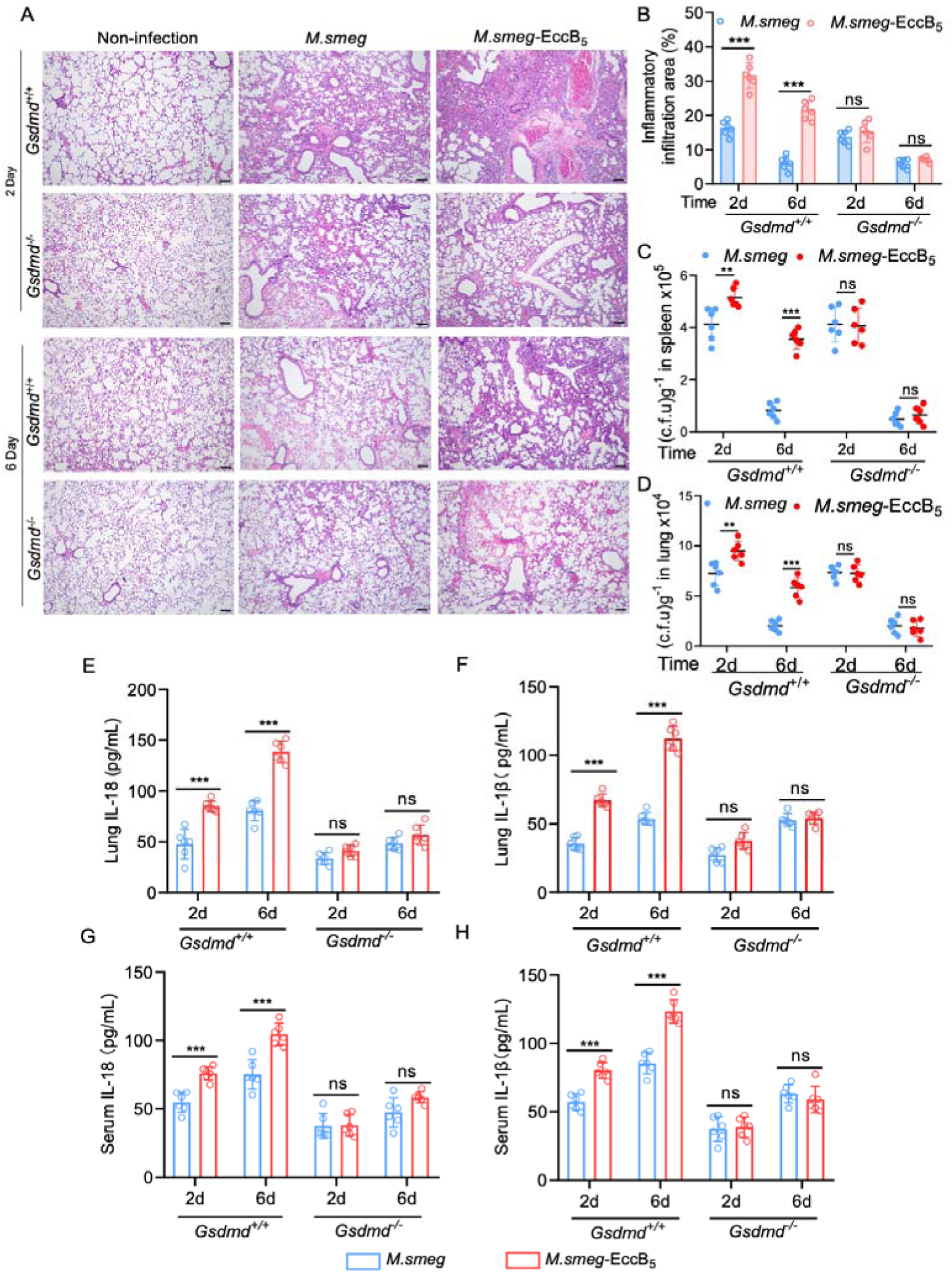
EccB_5_ induces pyroptosis to promote pathogen pathogenicity and dissemination in vivo. (**A**) Histopathology of lung sections from *Gsdmd*^+/+^ or *Gsdmd*^−/−^ mice (n=6 per group) intravenously infected with or without the indicated *M.smeg* or *M. smeg*-EccB_5_ for 2 to 6 days. Scale bars, 100 μm. (**B**) Quantification of inflammatory areas in lung sections from mice infected as in (**A**). **(C** and **D)** Bacterial colony-forming units (CFUs) recovered from lungs (**C**) and spleens (**D**) of mice infected as in (**A**). (**E** to **H**) ELISA analysis of cytokines (IL-1β and IL-18) in lung homogenates (upper panels) and sera (lower panels) from mice infected as in (**A**). Data are presented as mean ± SD of n = 6. Statistical significance was determined by two-way ANOVA followed by Tukey’s multiple comparisons test. ns, not significant (P > 0.05); P < 0.05; *P < 0.01; **P < 0.001; ***P < 0.0001.

## Discussion

Pathogenic microbes have evolved sophisticated strategies to evade and manipulate host immune defenses, enabling their persistence and systemic dissemination within the host [32, 33]. While prior studies have primarily focused on intracellular survival and immune evasion, the mechanisms by which pathogens promote their own spread remain incompletely understood. In this study, we employed an Mtb proteome microarray to identify mycobacterial proteins capable of interacting with human GSDMD (Fig. 1). The result shows that EccB_5_ directly binds to GSDMD (Fig. 2). Functional analyses demonstrated that EccB_5_ promotes caspase-1–mediated cleavage of GSDMD, thereby inducing pyroptosis both *in vitro* and *in vivo*, which in turn enhances Mtb pathogenicity and dissemination (Fig. 3-7).

EccB_5_ has previously been implicated in nutrient uptake, substrate transport, and bacterial viability [24, 34, 35]. Our findings extend its functional repertoire by implicating EccB_5_ in the modulation of host immune responses through direct engagement with pyroptotic signaling machinery. Mechanistically, we propose the formation of a tripartite complex comprising EccB_5_, caspase-1, and GSDMD, wherein EccB_5_ facilitates or stabilizes the interaction between caspase-1 and GSDMD, promoting efficient cleavage and activation of GSDMD (Fig. 6).

Pyroptosis is a form of GSDMD-mediated programmed inflammatory cell death [36–40], which is essential for eliminating infected cells and alerting neighboring immune cells, whereas excessive or dysregulated pyroptosis can contribute to immunopathology and facilitate pathogen dissemination [14, 18]. Recent studies have highlighted the importance of GSDMD-driven pyroptosis in controlling various intracellular infections, including those caused by *Salmonella* [41], *Shigella* [42], *Listeria* [43], *Bacillus* [44], and *Legionella* [45]. However, in the case of Mtb, pyroptosis has a particularly complex role [46, 47]. Pyroptotic cell death may contribute for bacterial egress, thereby facilitating spread within the host and to new hosts [48]. When host defenses fail to prevent potassium efflux and subsequent NLRP3 inflammasome activation, pyroptosis becomes inevitable, promoting Mtb release and propagation [18].

Our results position EccB_5_ as a key virulence determinant that modulates host cell death to the pathogen’s advantage. Mtb EccB_5_ directly targets GSDMD to drive pyroptosis, leading to enhanced release of inflammatory mediators and lung injury in infected mice. While this response contributes to initial bacterial control, it ultimately facilitates disease progression by promoting tissue damage and bacterial spread. These findings provide novel insight into the molecular mechanisms through which Mtb manipulates host pyroptotic pathways and highlight EccB_5_ as a potential target for host-directed therapeutic interventions.

### Methods summary

Full details of the materials and methods are provided in the supplementary materials. Briefly, all experiments were conducted in accordance with protocols approved by the institutional animal care and use committee.

For bacterial strain construction, Mtb H37Rv and its derivatives were cultured on Middlebrook agar or in broth, with genetic modifications introduced via homologous recombination. Plasmids were constructed and introduced into *E. coli* for cloning, and the resulting recombinant plasmids were used to transform *M. smegmatis*. Site-directed mutagenesis was employed to generate specific point mutations in Mtb genes. Lentiviral production and transfection were carried out in HEK 293T cells to generate stable cell lines for protein overexpression. In vitro assays were performed using RAW264.7, iBMDM, and THP-1 macrophage cell lines. Cell viability was measured with the CellTiter-Lumi™ assay, and cytotoxicity was assessed by lactate dehydrogenase (LDH) release. For immunofluorescence microscopy, cells were fixed, permeabilized, and stained with primary and secondary antibodies to visualize protein localization, with confocal imaging used to examine protein interactions. For infection assays, macrophages were infected with Mtb strains at a multiplicity of infection (MOI) of 10. After infection, cells were lysed for bacterial CFU counting, and cytokine levels were measured by ELISA. In vivo experiments were performed using C57BL/6 and *Gsdmd*^-/-^ mice infected with *M. smegmatis*, with tissues collected for histological analysis and CFU quantification. Protein purification was conducted using gel-filtration columns, followed by analysis with SDS-PAGE and western blotting. Immunoprecipitation was used to investigate protein-protein interactions. All experiments were repeated as indicated, and statistical analyses were performed using GraphPad Prism.

## Acknowledgements

We thank Dr. Zhaoyu Lin (Nanjing University) for providing the assistance with GSDMD knockout mice and Dr. Shengce Tao (Shanghai Jiaotong University) for supporting with the MTB microarray. This work was supported by grants from the National Natural Science Foundation of China (32161160323) and the Shanghai Committee of Science and Technology (24490713600).

## Authorship Contributions

J.L. conceived and designed the study. Y.S., Y.H., A.C., Y.L., Y.G., X.L., L.G., M.Y., Y.Q., L.Z., Y.S., and H.Y. performed the experiments and analyzed the data. Y.S. and J.L. analyzed the data and wrote the manuscript. All authors discussed the results and commented on the manuscript.

## Disclosure of Conflicts of Interest

The authors declare that they have no competing interests.

## Ethics Statement

All animal experiments were performed in accordance with the NIH Guide for the Care and Use of Laboratory Animals, with the approval of the Scientific Investigation Board of School of Life Sciences, Fudan University (2020-JS-016).

## Notes

### Competing Interest Statement

The authors have declared no competing interest.

