## Supplementary file for "The Mycobacterium tuberculosis ESX effector promotes pyroptosis-dependent pathogenicity and dissemination"

Yajie Shen *et al.*

**The PDF file includes:**

Materials and Methods

Figures. S1 to S7

**Other Supplementary Materials for this manuscript include the following:**

Tables S1 to S4

Materials and Methods

Bacterial strains and plasmids

Bacterial strains used in this study are described in table S3. *Mycobacterium tuberculosis* H37Rv strains were grown in Middlebrook 7H10 agar (BD Biosciences) supplemented with 10% oleic acid-albumin-dextrose-catalase (OADC) (BD Biosciences) or in Middlebrook 7H9 broth (BD Biosciences) supplemented with 10% OADC and 0.05% Tween-80 (Sigma). Mtb strains were cultured to mid-log phase (OD₆₀₀ ≈ 0.6) in the presence of 50 μg/mL kanamycin and either 100 or 200 ng/mL anhydrotetracycline (ATc). Bacterial titers (c.f.u./mL) were determined by serial dilution and plating on 7H10 agar, followed by incubation at 37 °C for 4 weeks. Aliquots of cultures were mixed with 20% glycerol in Middlebrook 7H9 medium and stored at –80 °C for subsequent use in macrophage and mouse infections, western blotting, and quantitative PCR analysis. *M. smegmatis* (strain ATCC 19420), *Escherichia coli* DH5α (strain ATCC 25922), and *E. coli* BL-21 (stain ATCC BAA-1025) were propagated from laboratory stocks (School of Life Sciences, Fudan University, Shanghai, China). *E. coli* DH5a and BL21 (DE3) were grown in LB medium. The PMV261 plasmid (kindly provided by Dr. Lu Zhang), was used to overexpress Mtb Rv1782c (EccB_5_) in *M. smegmatis*. Recombinant *M. smegmatis* expressing *M. tuberculosis* EccB_5_ (*M. smeg*-EccB_5_) was generated by electroporating pMV261-EccB_5_ into *M. smegmatis*. *M. smeg*-EccB_5_ were plated on 7H10 containing kanamycin (50 μg/mL). Transformants were selected on 7H10 agar supplemented with 50 μg/mL kanamycin and maintained in 7H9 medium containing the same antibiotic.

The full-length cDNA of EccB_5_ was amplified from Mtb H37Rv genomic DNA, and cloned into the pcDNA3.1-HA, pcDNA3.1-Flag, pCDH-GFP, pSMT3-His-SUMO, and PMV261 vectors, respectivly. PLJR965-EccB_5_ was a kind gift from Dr. Yi-Cheng Sun (Peking Union Medical College, Beijing, P. R. China.). pSMT3-GSDMD, HA-tagged GSDMD, mCherry-tagged GSDMD, and mutants of GSDMD were subcloned from pcDNA3.1-GSDMD-Flag as described previously. All primers used for cloning are listed in table S4. All plasmids were verified by DNA sequencing.

Construction of Mycobacteria strains

A CRISPR interference (CRISPRi) system was used to suppress Rv1782c expression in M. tuberculosis H37Rv. A specific single guide RNA (sgRNA; sequence: 5′- GACGCGGGTGGACTCTTTGG -3′), targeting the non-template strand of Rv1782c, was designed and cloned into the pLJR965 plasmid, which encodes a tetracycline-inducible dCas9, yielding pLJR965-sgEccB5. The recombinant plasmid was introduced into *M. tuberculosis* H37Rv via electroporation. Transformants were selected on 7H10 agar supplemented with 10% ADC and 50 μg/mL kanamycin at 37 °C. Kanamycin-resistant colonies were screened by PCR to confirm the integration of the dcas9 cassette, and knockdown efficiency was evaluated by assessing growth on ATc-containing 7H10 plates. Quantitative real-time PCR (qPCR) was used to assess the knockdown efficiency of Rv1782c, with gene expression normalized to the housekeeping gene sigA.

Mice

*Gsdmd*-knockout mice were kindly provided by Dr. Z. Lin (Nanjing University). Wild-type C57BL/6 mice were purchased from JieSiJie Biosciences (Shanghai, China). Female littermates (C57BL/6 background), 6–8 weeks of age, were used for all experiments. Mice were randomly assigned to experimental groups based on genotype. No blinding was performed, as data collection was conducted through objective and automated methods. All animal procedures were approved by the Scientific Investigation Board of the School of Life Sciences, Fudan University (protocol number: 2020-JS-016) and carried out in accordance with the NIH Guide for the Care and Use of Laboratory Animals.

Cell lines and culture conditions

The murine macrophage cell lines RAW264.7 and iBMDMs were cultured in Dulbecco’s Modified Eagle Medium (DMEM; Gibco) supplemented with 10% fetal bovine serum (FBS) and 100 µg/mL penicillin-streptomycin. The human THP-1 monocyte cell line was maintained in RPMI 1640 medium (Gibco) with 10% FBS and 100 µg/mL penicillin-streptomycin, seeded into 48-well plates at a density of 1 × 10⁵ cells per well, and differentiated into macrophages by treatment with 20 ng/mL phorbol 12-myristate 13-acetate (PMA) for 24 hours. All cells were maintained at 37 °C in a humidified incubator with 5% CO₂.

Cell transfection and lentiviral production

For transient transfection, HEK293T cells were transfected with plasmids using Lipofectamine 8000 (Beyotime, China) according to the manufacturer’s protocol. For stable expression, PCDH-based vector encoding EccB5 or empty control plasmid was co-transfected with lentiviral packaging plasmids psPAX2 and pMD2.G (Addgene, USA) into HEK293T cells using Lipofectamine 8000 at a ratio of 3:1:4 (vector: psPAX2: pMD2.G). Viral supernatants were collected at 48- and 72-hours post-transfection, filtered through a 0.45-μm membrane to remove cell debris, and concentrated using Lenti-X Concentrator (Takara, Japan) at a 4:1 ratio. Viral particles were pelleted by centrifugation at 1500 × g for 45 min at 4 °C and resuspended in antibiotic- and serum-free DMEM at 1/100 of the original culture volume.

Antibodies and reagents

Antibodies used in this study include mouse monoclonal anti-β-actin (Proteintech, 60008-1-Ig), mouse monoclonal anti-Flag M2 (Sigma-Aldrich, F1804), mouse polyclonal anti-HA (Proteintech, 66006-2-Ig), rabbit monoclonal anti-GFP (Proteintech, 66002-1-Ig), rabbit monoclonal anti-mCherry (Proteintech, 68088-1-Ig), rabbit polyclonal anti-GSDMD (Abcam, ab209845), rabbit polyclonal anti-cleaved N-terminal GSDMD (Abcam, ab215203), rabbit monoclonal anti-Cleaved caspase1 (Cell Signaling Technology, 89332), rabbit monoclonal anti-caspase1 (Cell Signaling Technology, 24232), rabbit monoclonal anti-Cleaved IL-1β (Cell Signaling Technology, 63124), rabbit monoclonal anti-IL-1β (Cell Signaling Technology, 31202), rabbit monoclonal anti-GSDME (Abcam, ab215191), rabbit monoclonal anti-caspase3 (Cell Signaling Technology, 9662), rabbit monoclonal anti-caspase4 (Cell Signaling Technology, 4450), rabbit monoclonal anti-caspase5 (Cell Signaling Technology, 46680), rabbit monoclonal anti-caspase11 (Cell Signaling Technology, 14340), rabbit lgG control antibody (Cell Signaling Technology, 2729), anti-Mycobacterium tuberculosis Ag85B (Abcam, ab312328), HRP-conjugated goat anti-mouse IgG (Proteintech , SA00001-1), HRP-conjugated goat anti‐rabbit IgG (Proteintech, SA00001-2), Alexa Fluor Plus 488-conjugated goat anti-mouse IgG (Invitrogen, A32723) and Alexa Fluor 549-conjugated goat anti-rabbit IgG (Invitrogen, A32740). Dimethyl sulfoxide (DMSO) and β-mercaptoethanol (2-ME) were purchased from Sigma-Aldrich. LPS (HY-D1056) and Nigericin (HY-127019) was purchased from MedChemExpress. The protease inhibitor PMSF tablets were from (Beyotime, ST506).

Recombinant protein purification

All procedures were performed at 4 °C unless otherwise stated. The *M. tuberculosis* EccB5 protein was expressed by transforming a plasmid encoding wild-type EccB5 into *E. coli* BL21 (DE3) cells. Cultures were grown in Luria–Bertani (LB) medium at 37 °C until the OD₆₀₀ reached 0.6–0.8, followed by induction with 0.2 mM isopropyl β-D-1-thiogalactopyranoside (IPTG) at 18 °C for 18–20 hours. Cells were harvested by centrifugation at 6000 rpm for 15 minutes, resuspended in lysis buffer (50 mM Tris-HCl pH 8.0, 300 mM NaCl, 5% glycerol) supplemented with 0.25% n-dodecyl-β-D-maltoside (DDM) and 6 mM MgCl₂, and lysed using a high-pressure homogenizer (JNBIO, China). Cell debris was removed by centrifugation at 17,000 rpm for 60 minutes, and the supernatant was loaded onto a HisTrap™ HP column (GE Healthcare). Bound proteins were eluted with buffer containing 50 mM Tris-HCl (pH 8.0), 500 mM NaCl, 500 mM imidazole, and 0.01% DDM. The His-SUMO tag was removed by overnight cleavage with ULP1 protease, followed by a second round of Ni-NTA affinity purification to remove uncleaved protein and the protease. The eluted protein was further purified by size-exclusion chromatography using a Superdex 200 16/600 column (GE Healthcare) equilibrated with buffer containing 20 mM Tris-HCl (pH 8.0) and 100 mM NaCl. Endotoxin removal was performed using polymyxin B-based endotoxin removal columns. The purity and integrity of the final protein preparation were verified by SDS–PAGE.

Mtb Proteome Microarray Screening

Recombinant human GSDMD protein with His tag (e.g., or Flag) was expressed in *E. coli* and purified by Ni-NTA affinity chromatography under native conditions. The Mtb proteome microarray, which contains over 4,000 individually expressed and spotted Mtb proteins, was pre-blocked with 5% BSA in PBST to reduce nonspecific binding. The purified GSDMD protein was then incubated with the microarray at 4°C overnight. After washing, binding events were detected using an anti-tag primary antibody and an appropriate secondary antibody conjugated to a fluorescent dye. Arrays were scanned using a microarray scanner (GenePix 4000B), and signal intensities were quantified with GenePix Pro software. Proteins with significantly higher fluorescence intensity compared to negative controls were considered putative GSDMD interactors and were selected for further validation.

Cell viability and Cytotoxicity Assay

Cell viability was determined by measuring ATP levels using the CellTiter-Lumi™ II Luminescent Cell Viability Assay Kit (Beyotime, C0056). Cell cytotoxicity of all the cells in a well was measured by the LDH release assay using LDH Cytotoxicity Assay Kit ((Beyotime, C0016)). To evaluate the loss of plasma membrane permeability and integrity, cells were cultured in medium supplemented with 2 μg/mL propidium iodide (Yeasen Biotechnology, 40711ES). The fluorescence intensity was continuously recorded upon the relevant treatment for 4 h with 5 min intervals using a BioTek Synergy H1 plate reader with Gene5 software.

Confocal microscopy

To assess the colocalization of EccB5 and GSDMD, HEK293T cells were seeded onto live-cell imaging dishes (Livefocus) and transfected with FLAG-EccB5 and HA-murine/human-GSDMD plasmids. After 36 hours, cells were washed with PBS, fixed with 4% paraformaldehyde at 4 °C, and permeabilized with 0.1% Triton X-100 in PBS. Blocking was performed with 3% bovine serum albumin (BSA; Sigma) in PBS for 1 hour. Cells were then incubated overnight at 4 °C with mouse anti-FLAG and rabbit anti-HA primary antibodies. After washing with PBS, cells were incubated with Alexa Fluor Plus 488-conjugated goat anti-mouse IgG and Alexa Fluor 549-conjugated goat anti-rabbit IgG for 1 hour at room temperature. Nuclei were counterstained with DAPI (Beyotime, C1002) for 10 minutes. Colocalization images were obtained using a laser scanning confocal microscope (Zeiss-LSM880).

ELISA for cytokine measurement

IL‐1β or IL-18 expression levels in supernatants were analyzed by an ELISA kit (Multi sciences, EK218, EK201B) following the manufacturer's instructions.

Protein electroporation

Recombinant full-length EccB_5_ was electroporated into iBMDMs or *Gsdmd*^-/-^ iBMDMs cells, respectively, to test their pore-forming and pyroptosis-inducing activities. Protein electroporation was performed using the Neon™ Electroporation System (Thermo Scientific). Briefly, 2 × 10^6^ cells were washed with PBS (without Ca^2+^ and Mg^2+^) by centrifugation at 400 g for 5 minutes at room temperature. Aspirate the PBS and resuspend the cell pellet in Resuspension Buffer R, and electroporated using the following parameters (1900 V, 30 ms and 1 pulse). Cell viability was measured by CellTiter-Lumi™ 2 h after electroporation.

Macrophages infection

Macrophages were seeded and cultured overnight before infection with *M. tuberculosis* H37Rv, H37Rv EccB5_cKDTet, *M. smegmatis*, or M. smeg-EccB5 strains at a multiplicity of infection (MOI) of 10. At indicated time points post-infection, supernatants were collected for lactate dehydrogenase (LDH) release and inflammatory cytokine assays, while cell lysates were harvested for Western blot analysis. For bacterial survival assays, wild-type (WT) and *Gsdmd*⁻/⁻ iBMDMs were infected with *M. smegmatis* strains at MOI = 10 for 2 hours in DMEM. After infection, the medium was removed, and cells were washed three times with PBS to remove extracellular bacteria. Cells were then incubated in fresh DMEM containing 10 μg/mL gentamicin for an additional period to eliminate any remaining extracellular bacteria. After incubation, cells were washed again three times with PBS and lysed in 7H9 medium containing 0.05% SDS for 10 minutes. Serial 10-fold dilutions of the lysates were prepared in 0.05% Tween-80, and aliquots were plated on 7H10 agar plates to determine colony-forming units (CFUs).

Immunoblotting analysis and immunoprecipitation

Cells were washed twice with phosphate-buffered saline (PBS) and subsequently lysed in radioimmunoprecipitation assay (RIPA) buffer on ice for 60 minutes. The lysis buffer contained 20 mM Tris-HCl (pH 7.5), 150 mM NaCl, 1% Triton X-100, 1 mM phenylmethylsulfonyl fluoride (PMSF), and 2 mM ethylenediaminetetraacetic acid (EDTA). Following incubation, the lysates were clarified by centrifugation at 16,200 g for 10 minutes at 4°C. Protein concentration was determined using the BCA protein assay kit (Beyotime, China). Equal amounts of protein (typically 30–50 μg) were separated by SDS-PAGE on a 10% acrylamide gel and transferred onto polyvinylidene difluoride (PVDF) membranes (Millipore, USA) according to the standard protocol. To assess protein-protein interactions, co-immunoprecipitation (Co-IP) assays were performed. Protein lysates were incubated with magnetic beads pre-conjugated with specific antibodies for 2 hours at room temperature. The Co-IP magnetic beads (HY-K0202, MCE, USA) were prepared according to the manufacturer’s instructions. After incubation with the antibodies, the protein-bead complexes were mixed with the prepared lysates and incubated for an additional 2 hours. The magnetic beads were then washed three times with PBS, followed by elution in a buffer containing loading dye. The eluted samples were analyzed by SDS-PAGE, and the proteins were transferred onto PVDF membranes. After transfer, the membranes were blocked for 1 hour at room temperature using 5% non-fat dry milk or bovine serum albumin (BSA) in tris-buffered saline (TBS). Membranes were then incubated overnight at 4°C with primary antibodies diluted in TBS containing 0.5% Tween 20 (TBST). After primary antibody incubation, membranes were washed three times with TBST and incubated with a horseradish peroxidase (HRP)-conjugated secondary antibody (1:10,000, Cell Signaling Technology) for 1 hour at room temperature. The protein bands were visualized by enhanced chemiluminescence (ECL, Tanon, 180-5001) and imaged using a gel documentation system. Quantification of Western blotting bands was performed using ImageJ software, and statistical analysis was conducted to compare the expression levels across experimental groups.

Histological analysis

Lungs and livers from wild-type (WT) *M. smegmatis* (M. smeg) and M. smeg-EccB5-infected or control mice were harvested, fixed in 4% paraformaldehyde, and embedded in paraffin. The tissues were then cut into sections and mounted on adhesive microscope slides. Hematoxylin and eosin (H&E) staining was performed following standard protocols to examine tissue morphology. The stained sections were imaged using light microscopy (Olympus BX53) and analyzed with CellSens software.

Mice infection

For the *M. smegmatis* challenge model, 6- to 8-week-old female C57BL/6 mice were intravenously infected with either WT *M. smeg* or *M. smeg*-EccB5 at a dose of 10^8^ CFU per mouse. Briefly, bacterial cultures in the logarithmic growth phase were washed twice with 1× PBS, sonicated to create a single-cell suspension, and adjusted to an OD600 of 0.1 in 1× PBS. The suspension was then used for intravenous infection. Mice were euthanized at the designated time points. Lung and spleen tissues were harvested, fixed in 4% paraformaldehyde, and subjected to H&E staining for histopathological analysis. Lung and spleen homogenates were used for qPCR analysis and colony-forming unit (CFU) counting. Serum cytokine levels were measured by ELISA. All mice were housed in microisolator cages, provided with standard laboratory diet and water, and maintained at the animal facility of Fudan University. Animal experiments were conducted in accordance with the National Institutes of Health (NIH) Guide for the Care and Use of Laboratory Animals and approved by the Scientific Investigation Board of the School of Life Sciences, Fudan University (2020-JS-016).

Statistical analyses

Data were shown as means ± SD of three technical replicates. All data met the assumptions of the tests. Student’s unpaired t test was used to compare the means of the two groups. Two-way analysis of variance (ANOVA) with Bonferroni posttests was used to compare the means between multiple groups. Data were analyzed by GraphPad Prism 9.0. P‐values < 0.05 were termed as significant (*P < 0.05; **P < 0.01; and ***P < 0.001); ns means non‐significant where P ≥ 0.05

**Supplementary Figures and Legends**

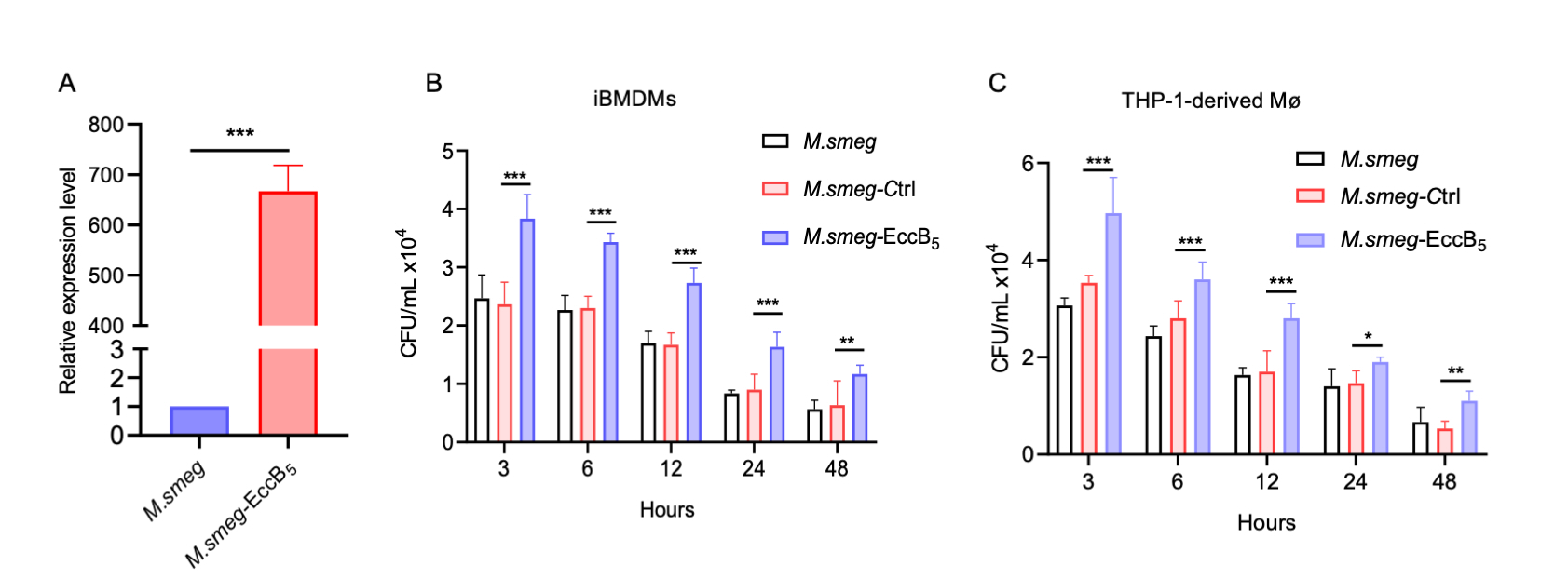
 **Fig. S1. Construction of *M. smeg* and *M. smeg*-EccB_5_. (A)** Relative expression level of EccB_5_ in *M. smeg* and *M. smeg*–EccB_5_ strain, respectively. (**B-C**) Bacterial intracellular survival analysis iBMDM and THP1 cells upon infection with *M. smeg*, *M. smeg* -control, and *M. smeg*–EccB_5_ strains.

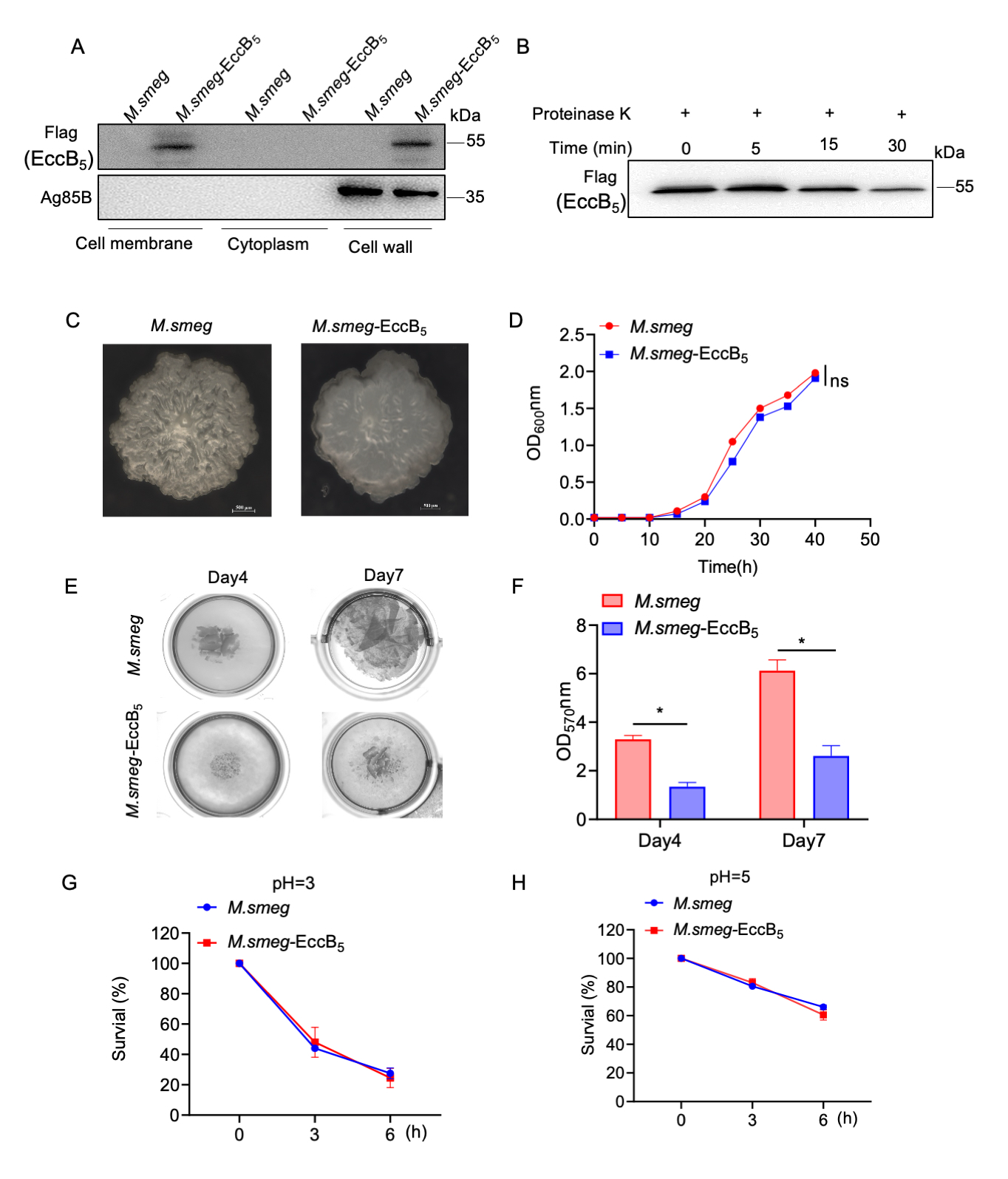
 **Fig. S2. Phenotype characterization of *M. smeg* and *M. smeg*-EccB_5._ (A**) Cell fractionation experiments were performed to determine the sub-cellular localization of EccB_5_, Ag85B protein serves as a cell wall marker of *M. smegmatis*. **(B**) Proteinase K treatment of M. smegmatis–EccB_5_ strains for the indicated times, followed by immunoblotting with anti-FLAG antibody. (**C**) The colony morphology of the recombinant strain in 7H10 medium. (**D**) The growth of recombinant *M. smeg* or *M. smeg*–EccB_5_ strains. (**E**) Cultures were incubated under stationary conditions at 37 °C in 48-well plates to induce Biofilm formation, and morphological changes were recorded by microscopy. (**F**) Biofilm formation was quantified by crystal violet staining. (**G** and **H**) In vitro growth of recombinan***t*** *M. smeg* and *M. smeg*-EccB_5_ after treatment with different pH gradient. Data are presented as mean ± SD of three independent groups. Statistical significance was determined by two-way ANOVA followed by Tukey’s multiple comparisons test. ns, not significant (P > 0.05); P < 0.05; *P < 0.01; **P < 0.001; ***P < 0.0001.

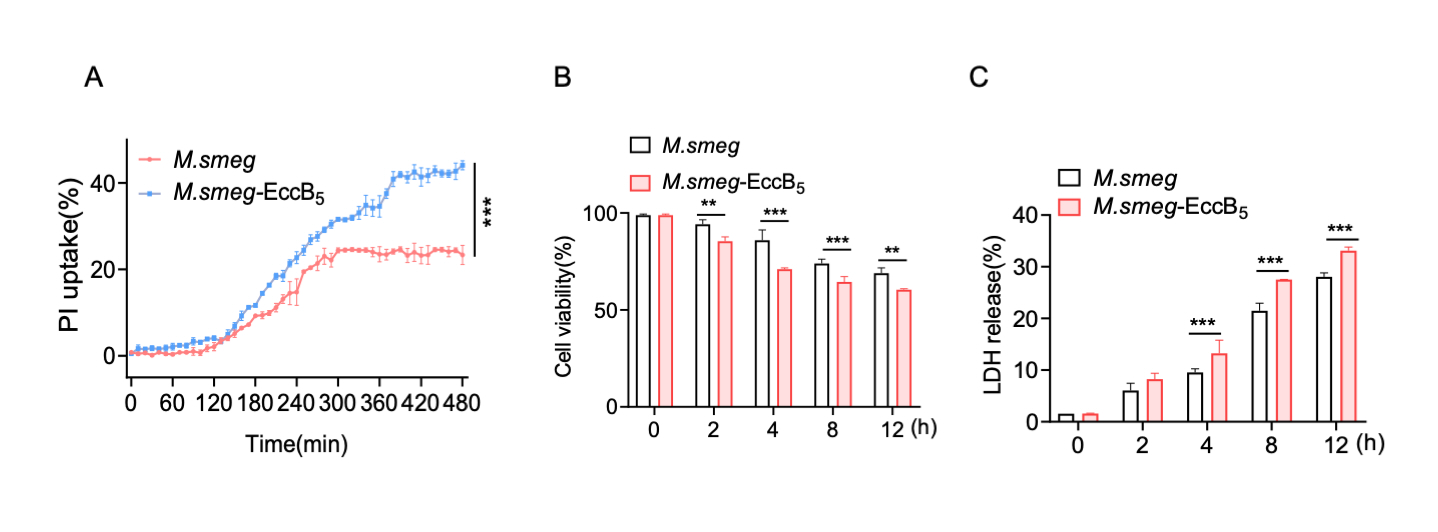
 **Fig. S3. Overexpression EccB_5_ in *M. smeg* induces pyroptosis of host cells.** iBMDM cells infected with *M. smeg* or *M. smeg*–EccB_5_, respectively, were detected for cytotoxicity by PI uptake (**A**), cell viability (**B**), or LDH release assays (**C**).

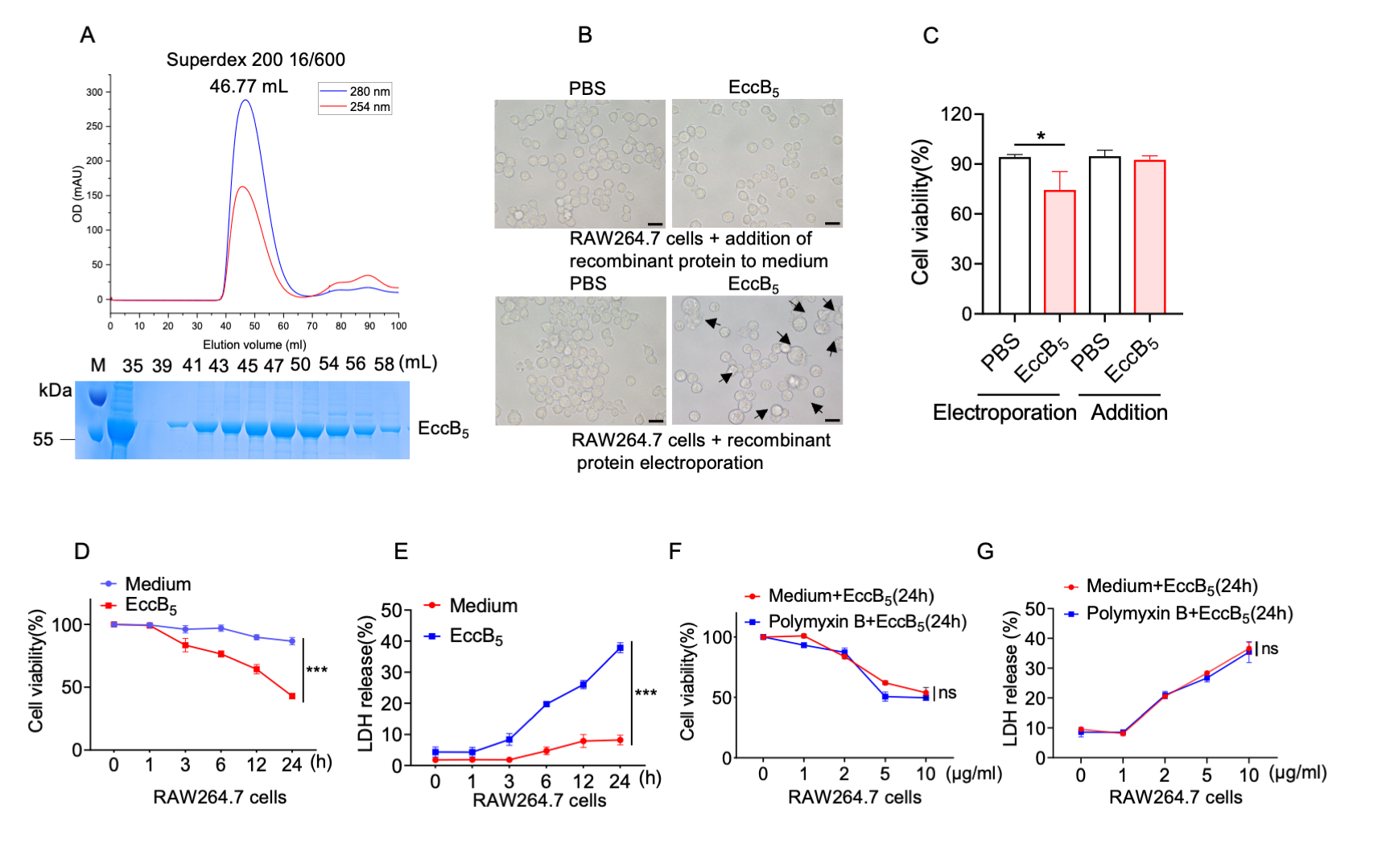
 **Fig. S4.** **Mtb EccB_5_ induces pyroptosis of host cells.** (**A**) Size-exclusion chromatographic profile and SDS-PAGE result of the EccB_5_ protein on a Superdex200 16/600 GL column. M: protein marker. (**B-C**) The recombinant EccB_5_ protein was introduced into RAW264.7 cells via electroporation or direct addition to the culture medium. The cell morphologies were observed, and the cell viability was analyzed. Scale bars, 20 μm. **(D-E)** Recombinant WT EccB_5_ or not infected (PBS) were electroporated into RAW264.7 cells for assessed cytotoxicity by cell viability (**D**) and LDH release assay (**E**). (**F-G**) Peritoneal macrophages pretreated with Poly. B followed by EccB_5_ treatment. Cell viability (**F**) and LDH release (**F**) were detected. Data are presented as mean ± SD of three independent groups. Statistical significance was determined by two-way ANOVA followed by Tukey’s multiple comparisons test. ns, not significant (P > 0.05); P < 0.05; *P < 0.01; **P < 0.001; ***P < 0.0001.

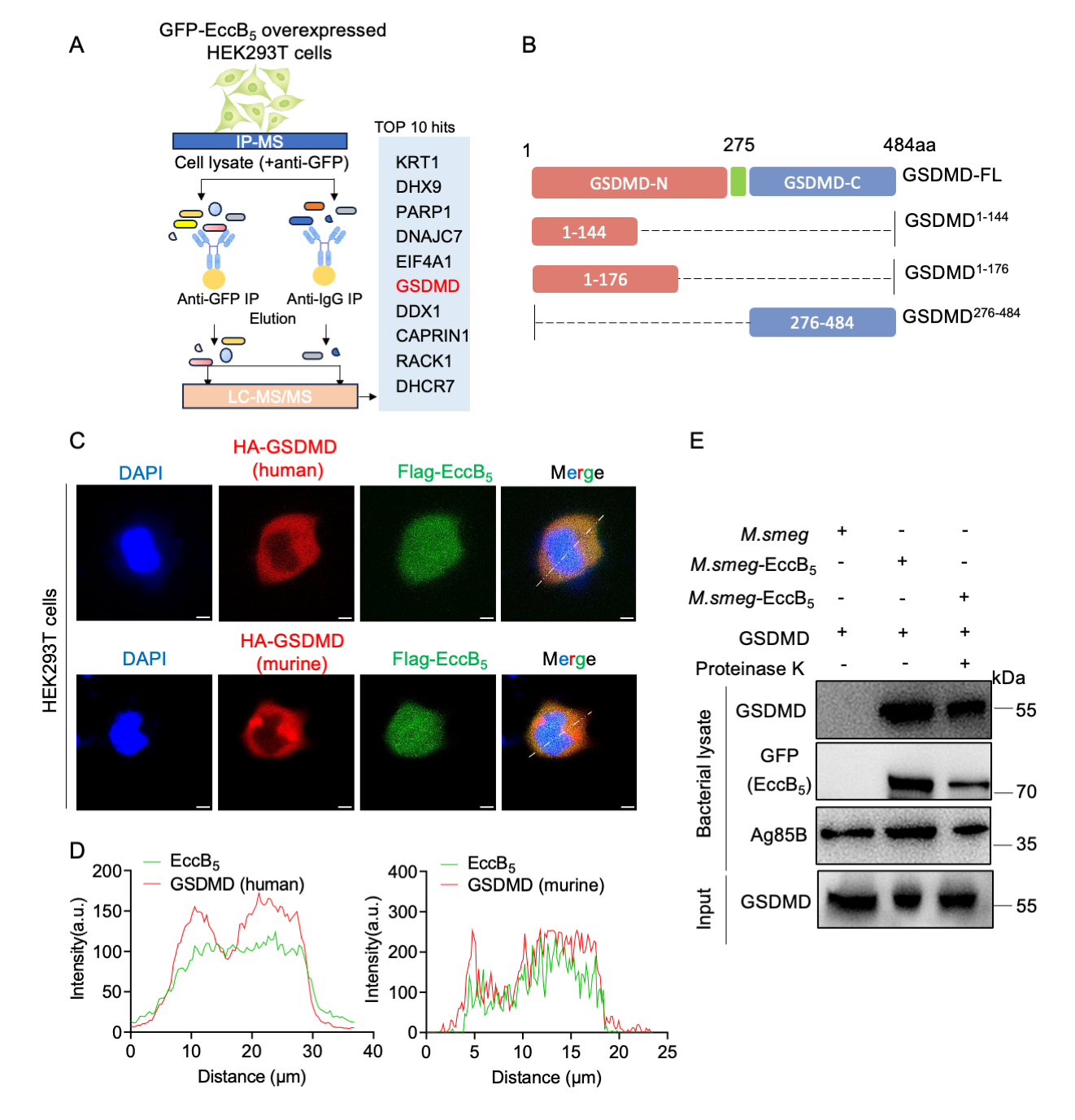
 **Fig. S5. Mtb EccB_5_ binds directly to GSDMD.** (**A**) Schematic of the IP–MS approach. (**B**) The schematic diagram of human GSDMD and its truncated mutants. (**C**) HEK293T cells were transfected with HA-human-GSDMD plasmid or HA-murine-GSDMD plasmid along with Flag-EccB_5_ plasmid. The cells were processed for confocal microscopy with anti-Flag antibody (EccB_5_) and anti-HA antibody (GSDMD). Nuclei were stained with DAPI. Scale bars, 5 μm. **(D)** Normalized intensity profiles are drawn from the white lines in (**D**) and show the relative pixel intensity along the line with regards to the distance and fluorescence wavelength (red, Cy3; green, Alexa 488). **(E)** Immunoblot analysis of whole bacterial lysate of Mtb cells pretreated with (+) or without (-) proteinase K followed by incubation with GSDMD at 4 °C for 4 h.

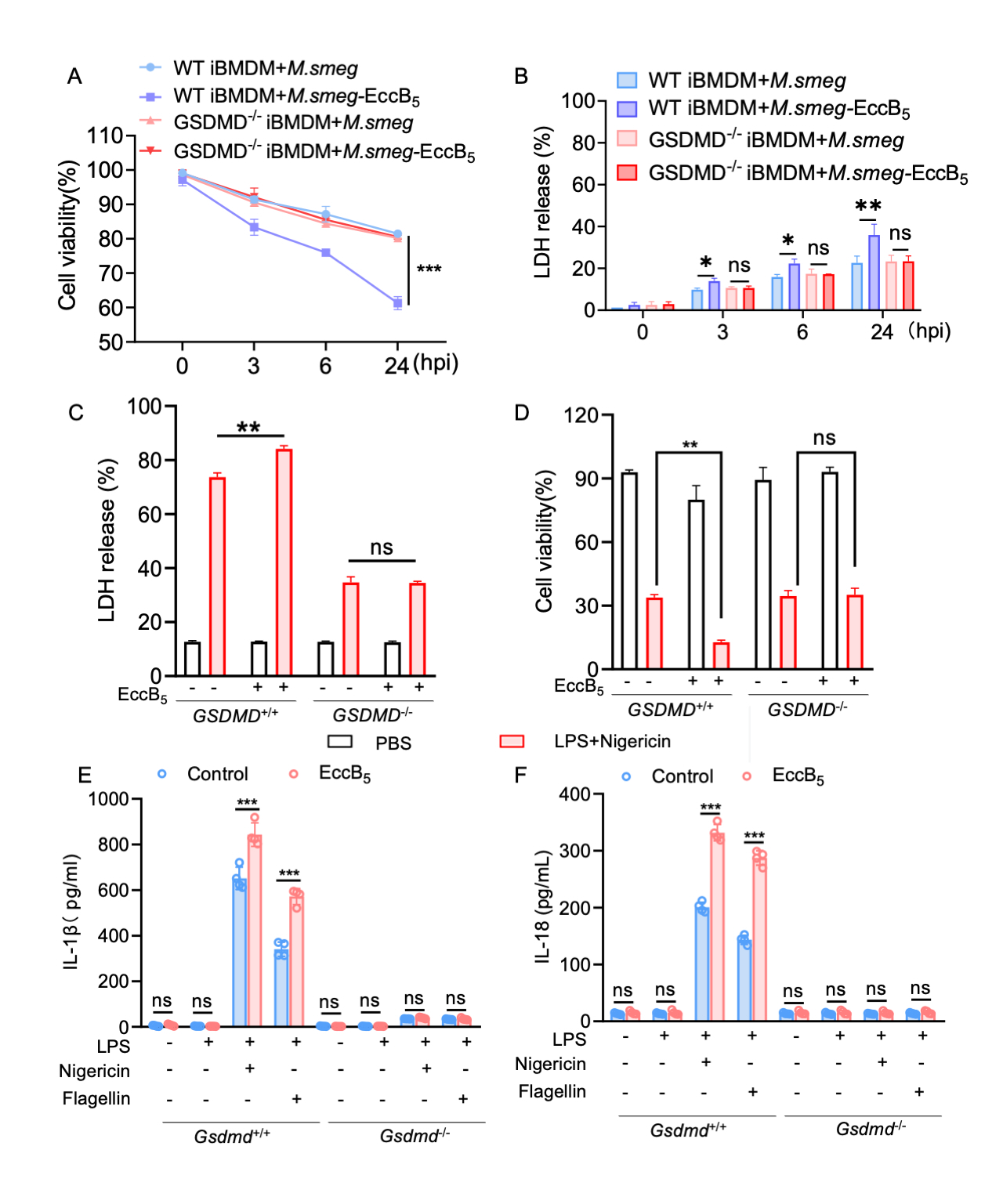
 **Fig. S6. EccB_5_ triggers pyroptotic cell death via GSDMD (A**-**B)** WT and *Gsdmd*^-/-^ iBMDM cells infected with *M. smeg* or *M. smeg*–EccB_5_ were evaluated by cell viability (**A**) and LDH (**B**). **(C-F)** WT and *Gsdmd*^-/-^ iBMDM cells stably expressing the EccB_5_ gene or control vector were stimulated with 10 μM nigericin for 30 min following pretreatment with 1 μg/ml LPS for 3 h. The percentage of cells undergoing pyroptosis was evaluated by LDH release (**C**), cell viability (**D**), and ELISA analysis for IL-1β and IL-18 (**E-F**). Data are presented as mean ± SD of three independent groups. Statistical significance was determined by two-way ANOVA followed by Tukey’s multiple comparisons test. ns, not significant (P > 0.05); P < 0.05; *P < 0.01; **P < 0.001; ***P < 0.0001.

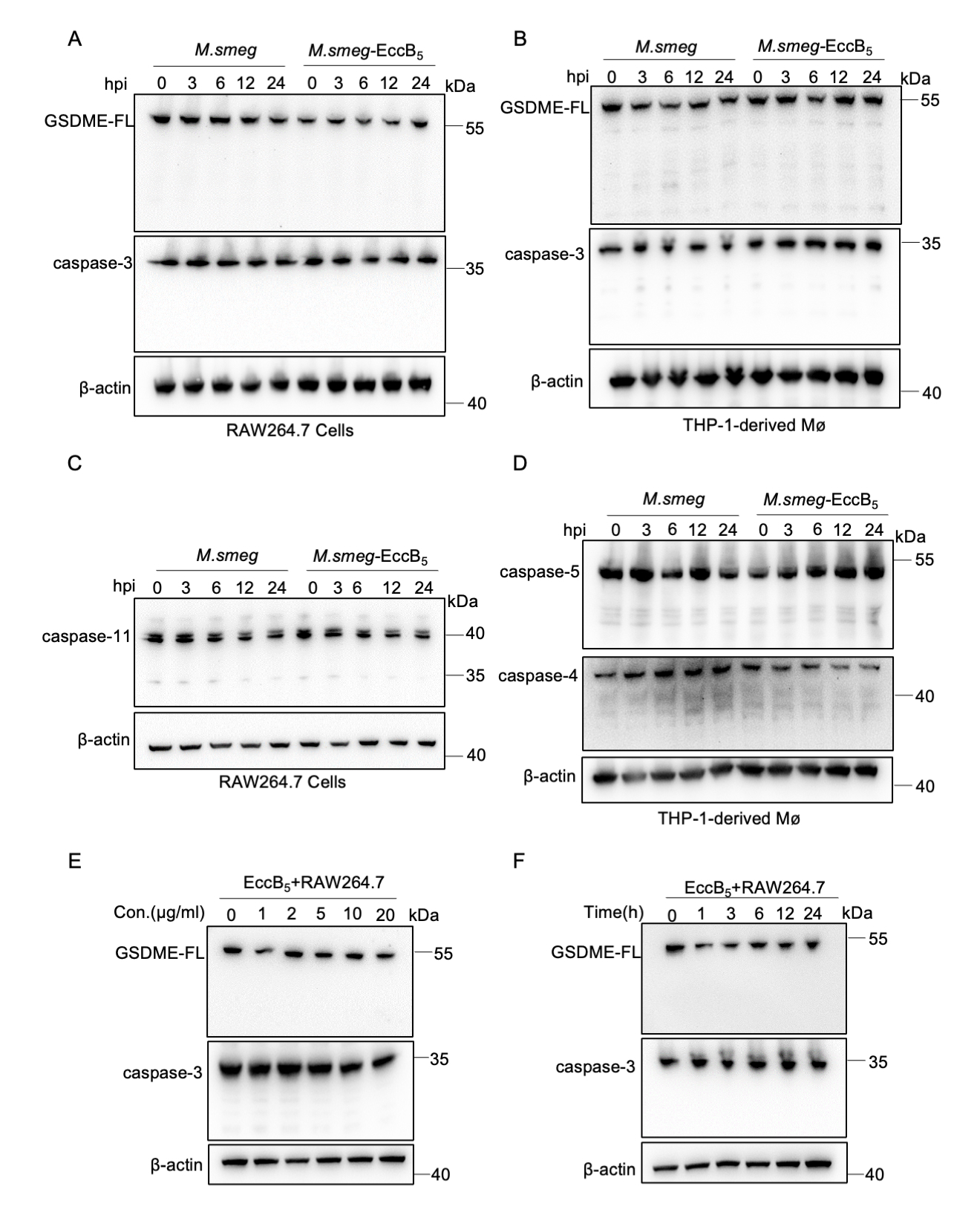
 **Fig. S7. EccB_5_ does not activate caspase-3/GSDME or caspase-4/5/11 signaling in macrophages.** (**A**-**D**) RAW264.7 cells (**A, C**) or THP-1-derived macrophages (**B, D**) infected with *M. smeg* or *M. smeg*–EccB_5_ were detected for caspase-3/GSDME and caspase-4/5/11 activation by WB. (**E** and **F**) RAW264.7 cells were electroporated with recombinant EccB_5_ protein at various concentrations (**E**) or electroporated with 2 μg/mL EccB_5_ for different durations (**F**) as indicated. Cell pellets were collected and immunoblotted with antibodies against the indicated proteins.
